# Variation in temperature of peak trait performance constrains adaptation of arthropod populations to climatic warming

**DOI:** 10.1101/2023.01.18.524448

**Authors:** Samraat Pawar, Paul J. Huxley, Thomas R. C. Smallwood, Miles L. Nesbit, Alex H. H. Chan, Marta S. Shocket, Leah R. Johnson, Dimitrios - Georgios Kontopoulos, Lauren Cator

## Abstract

The capacity of arthropod populations to adapt to long-term climatic warming is uncertain. Here, we combine theory and extensive data on diverse arthropod taxa to show that their rate of thermal adaptation to climatic warming will be constrained in two fundamental ways. First, the rate of thermal adaptation is predicted to be limited by the rate of shift in the temperature of peak performance of four life-history traits in a specific order: juvenile development, adult fecundity, juvenile mortality, and adult mortality. Second, thermal adaptation will be constrained due to differences in the temperature of peak performance among these four traits, which are expected to persist because of trade-offs. By compiling a new global dataset of 61 diverse arthropod species, we find strong evidence that contemporary populations have indeed evolved under these constraints. Our results provide a basis for using relatively feasible trait measurements to predict the adaptive capacity of diverse arthropod populations to climatic warming.

## Introduction

Arthropods are highly diverse and constitute almost half of the biomass of all animals on earth, fulfilling critical roles as prey, predators, decomposers, pollinators, pests, and disease vectors in virtually every ecosystem ^1^. Arthropod populations are under severe pressure globally due to pollution and land use changes, which will likely be compounded by ongoing and future climate change ^2,3,4,5,6^. The ability of their populations to adapt to climatic warming in particular has far-reaching implications for ecosystem functioning, agriculture, and human health ^7,8,9^.

The ability of a population to persist—its fitness—depends on its maximal (or “intrinsic”) growth rate (*r*_*m*_) in a given set of conditions. The response of arthropod *r*_*m*_ to environmental temperature is unimodal, with its peak typically occurring at a temperature (*T*_*opt*_) closer to the upper, rather than to the lower lethal limit (a left-skewed temperature-dependence; Fig. 1A) ^10,11,12^. This temperature-dependence of *r*_*m*_ emerges from the thermal performance curves (TPCs) of underlying life history traits (Fig. 1B) ^13,14,15^. Previous work has focused on the effect of thermal sensitivity (“*E*” in Fig. 1B) ^14,16^ or upper lethal thermal limit (*CT*_*max*_) ^17,18,19^ of traits on the temperature-dependence of *r*_*m*_, providing insights into the responses of populations to short-term thermal fluctuations and heat waves ^13,20,21^. However, to understand how populations will respond to long-term sustained climatic warming, we need to quantify the adaptive potential of *T*_*opt*_ and the corresponding *r*_*m*_ at that temperature (i.e., *r*_*opt*_; Fig. 1A), a measure of (thermal) fitness. Adaptive shifts in *T*_*opt*_ are primarily governed by shifts in the *T*_*pk*_s of underlying traits (Fig. 1B) ^14,15^. These trait-specific *T*_*pk*_s can evolve relatively rapidly under selection because they are subject to weaker thermodynamic constraints than thermal sensitivity (*E*) or *CT*_*max*_ ^12,22,23^. However, arthropods vary considerably in the form of their complex, stage-specific life histories, and a general, mechanistic trait TPC-based approach to quantify their adaptive potential to climate change has proven challenging ^9,21,24,25^.

**Figure 1:**
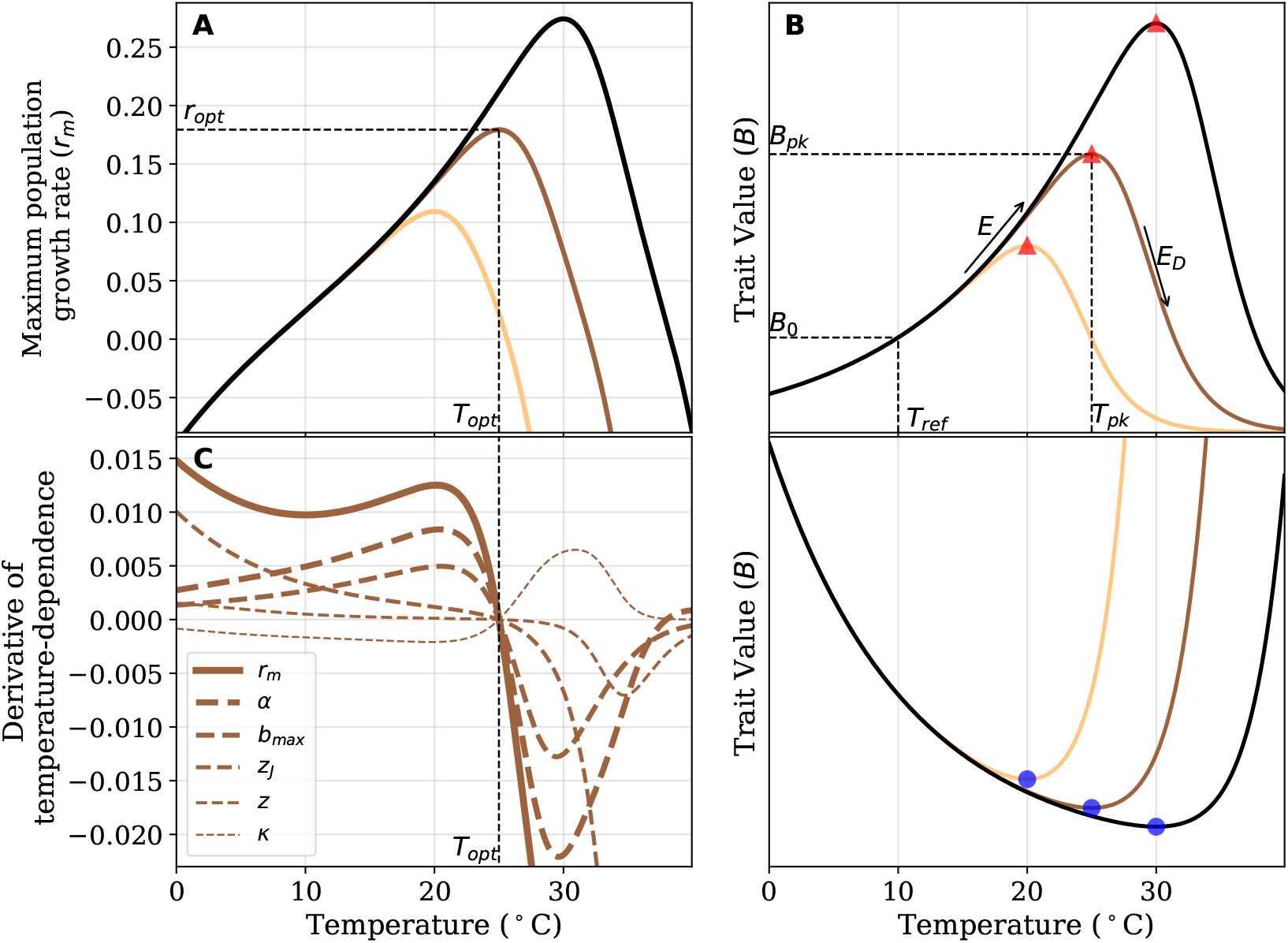
The relationship between the temperature-dependence of population fitness and its underlying traits. In all plots, the three curves represent populations adapted to three different temperatures. **A**: The temperature-dependence of population growth rate *r*_*m*_ (Equation (2)). Thermal fitness is the peak value that *r*_*m*_ reaches (*r*_*opt*_) and *T*_*opt*_ is the temperature at which this peak is achieved. **B:** *(Upper)* Illustration of the underlying trait TPCs modeled using the Sharpe-Schoolfield equation (Methods, Equation (3) and Table 1)), appropriate for development rate 1*/α*, maximum fecundity *b*_*max*_, and fecundity decline rate *κ*. **B:** *(Lower)* Illustration of a trait TPC modeled using the inverse of the Sharpe-Schoolfield equation, appropriate for juvenile mortality rate *z*_*J*_ and adult mortality rate *z*. Both upper and lower sets of curves were generated with arbitrary values for TPC parameters chosen for illustration from empirically reasonable ranges (Methods, Table 1). The y-axes values are not shown to keep focus on the qualitative shapes of these TPCs. **C**: Relative contributions of trait TPCs to the temperature dependence of *r*_*m*_. The greater the distance between a trait’s partial derivative (dashed curves, i.e., 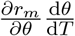where *θ* denotes any one of the five traits) and the total derivative 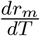 (solid curve), the smaller its contribution to *r*_*m*_’s temperature dependence.

Here we combine metabolic and life history theories to link variation in trait *T*_*pk*_s to thermal fitness and adaptive potential in the face of long-term directional changes in temperature (e.g., climatic warming), across diverse arthropod populations. Our approach simplifies the diverse complex stage-structures seen in arthropods to the temperature-dependence of five life-history traits, allowing general predictions that can be applied across taxa. We test our predictions with a global data synthesis of a diverse set of 61 arthropod taxa to reveal two fundamental constraints on thermal adaptation across arthropod lineages.

**Table 1:** Definitions of model parameters.

| Parameter | Units | Description |
| --- | --- | --- |
| $r_m$ | $\text{day}^{-1}$ | Maximal population growth rate |
| $\alpha$ | days | Egg to adult development time |
| $b_{max}$ | $\text{eggs} \times (\text{female} \times \text{day})^{-1}$ | Maximum fecundity rate |
| $\kappa$ | $\text{day}^{-1}$ | Fecundity loss rate |
| $z$ | $\text{day}^{-1}$ | Adult mortality rate |
| $z_J$ | $\text{day}^{-1}$ | Mortality rate averaged across juvenile stages |
| $B$ | measurement unit of trait | Trait value at a given temperature |
| $B_0$ | measurement unit of trait | Normalisation constant for trait value at $T_{ref}$ |
| $T_{pk}$ | $^{\circ}\text{C}$ or $\text{K}$ | Temperature at which trait performance peaks |
| $B_{pk}$ | Measurement unit of trait | Trait performance achieved at $T_{pk}$ |
| $T_{opt}$ | $^{\circ}\text{C}$ or $\text{K}$ | Temperature at which $r_m$ peaks |
| $r_{opt}$ | $\text{day}^{-1}$ | $r_m$ achieved at $T_{opt}$ |
| $E$ | eV | Activation energy, also called thermal sensitivity |
| $E_D$ | eV | Deactivation energy |
| $k$ | $\text{eV } K^{-1}$ | Universal Boltzmann constant ($8.617 \cdot 10^{-5}$) |

## Results

### The trait-driven temperature-dependence of fitness

We start with a mathematical equation for the temperature-dependence of *r*_*m*_ (Fig. 1A; Methods, Equation (2)) as a function of the TPCs for five key life history traits (Fig. 1) ^15^: juvenile-to-adult development time, *α*; juvenile mortality rate, *z*_*J*_ ; maximum fecundity, *b*_*max*_; fecundity decline with age, *κ*; and adult mortality rate, *z*. This equation predicts that the temperature-dependence of *r*_*m*_ increases as a population adapts to warmer temperatures (i.e., thermal fitness rises with *T*_*opt*_; Fig. 1A). This “hotter-is-better” pattern is consistent with the findings of empirical studies on arthropods as well as other ectotherms ^11,26^. In our theory, this pattern arises through thermodynamic constraints on life history traits built into the Sharpe-Schoolfield equation for TPCs (Methods, Equation (3)), which focuses on a single, rate-limiting enzyme underlying metabolic rate ^27^. While previous theoretical work has sought to understand the evolutionary basis and consequences of the hotter-is-better phenomenon ^11,17,21,23,26,28^, to the best of our knowledge, this is the first time that the contributions of underlying traits to it have been quantified. We note that our results do not rely on the the specific form or underlying thermodynamic assumptions of the Sharpe-Schoolfield equation; as such, any TPC model that encodes the hotter-is-better pattern, commonly observed across biological traits (Supplementary Results Section 1.1; ^14^), will yield qualitatively the same results.

### The existence of a hierarchy of traits driving thermal fitness

We next performed a trait sensitivity analysis to dissect how trait TPCs shape the temperature-dependence of *r*_*m*_ (Fig. 1C). It shows that populations will grow (*r*_*m*_ will be positive) as long as the negative fitness impact of an increase in juvenile and adult mortality rate (*z*_*j*_ and *z*) with temperature is counteracted by an increase in development and maximum fecundity rate (*α* and *b*_*max*_). So, for example, *r*_*m*_ rises above 0 at *∼*9^*°*^C (Fig. 1A) because at this point *z*_*J*_ and *z* fall below, and *α* and *b*_*max*_ rise above, a particular threshold (Fig. 1C). Similarly, the decline of *r*_*m*_ beyond *T*_*opt*_ is determined by how rapidly each of the underlying trait values change with temperature beyond that point (determined by their respective *E*_*D*_s). More crucially, this trait sensitivity analysis reveals a hierarchy in the importance of life history traits in driving both the location of thermal fitness along the temperature gradient (i.e., *T*_*opt*_) and its height (i.e., the thermal fitness achieved) (Fig. 1C): the TPC of development time (*α*) has the greatest influence followed by maximum fecundity (*b*_*max*_), juvenile mortality (*z*_*J*_), adult mortality (*z*) and fecundity loss rate (*κ*), with each of the latter three having a particularly weak effect *∼*5^*°*^C around *T*_*opt*_ (Fig. 1C). It is only at extreme temperatures (Fig. 1C, *T <* 10^*°*^C and *T >* 35^*°*^C) that juvenile mortality (*z*_*j*_) in particular exerts a strong influence on the temperature-dependence of *r*_*m*_. Thus the influence of the five traits in the region of thermal fitness can be ordered as *α > b*_*max*_ *> z*_*J*_ *> z > κ*. This hierarchy is expected to be a general result, consistent with the type of life stage structure typical of arthropod, and especially insect, species where the juvenile stages altogether are both more abundant and longer-lived than adults ^13,21^.

### The trait hierarchy shapes thermal fitness

Next we focused on trait *T*_*pk*_s as key targets of selection leading to thermal adaptation, that is, maximization of thermal fitness in a given constant thermal environment. We calculated thermal selection gradients and strengths of selection for each trait’s *T*_*pk*_ (Fig. 2). For the range of environmental temperatures we consider (0^*°*^C to 35^*°*^C), *T*_*opt*_ varies by about 12^*°*^C (*∼*16^*°*^C to 28^*°*^C). Every trait’s thermal selection gradient is necessarily asymptotic (Fig. 2E-H) because, even if the *T*_*pk*_ of that trait keeps increasing with environmental temperature, all the other traits still decline with temperature beyond their respective *T*_*pk*_ (Fig. 1B). These calculations predict that selection is strongest on the temperature of peak performance of development time (*α*), followed by those of the other traits in the same order 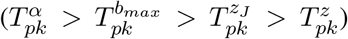, in line with our trait sensitivity analysis (Fig. 1C). We excluded *κ* from this analysis because it has a very weak influence on *r*_*m*_ (Fig. 1C), and TPC data are also lacking for it (Methods). Supplementary Results Section 1.7 shows that our results are indeed robust to variation in *κ*’s temperature dependence.

**Figure 2:**
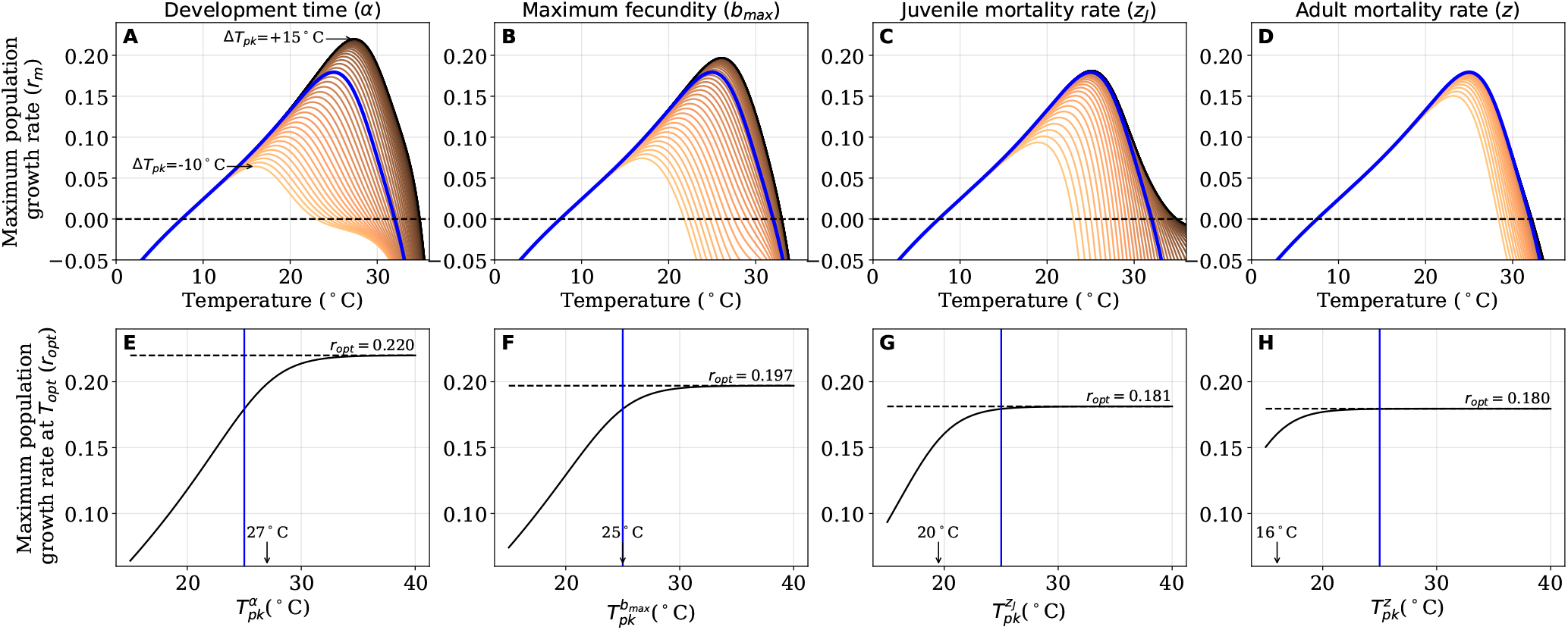
Thermal selection gradients for four key arthropod life-history traits. **A-D:** Change in the TPC of *r*_*m*_ with changes in *T*_*pk*_s of each of the four traits relative to all the others (Δ*T*_*pk*_). Δ*T*_*pk*_ ranges from *−*10^*°*^C (light lines) to +15^*°*^C (dark lines), with the solid blue line indicating the scenario in which all traits peak at the same temperature (Δ*T*_*pk*_ = 0) **E-H**: The corresponding selection gradients for each trait (Methods). The solid black line represents *r*_*m*_. The solid blue line represents Δ*T*_*pk*_ = 0; any increase (decrease) in *r*_*m*_ to the right (left) of this point represents potential increase (decrease) in thermal fitness that could be gained (lost) by increasing (decreasing) the *T*_*pk*_ of that trait relative to others. Both sets of plots (A-D & E-H) are ordered by decreasing strength of selection on the trait, that is, by the temperature at which the selection gradient begins to asymptote (vertical arrows in E-H) (Methods). The vertical blue lines in E-H again mark where Δ*T*_*pk*_ = 0.

These predictions provide a theoretical explanation for previous empirical observations about the importance of development rate, relative to other traits, for ectotherm thermal fitness ^29,30^, as well as its propensity to peak at higher temperatures relative to other traits in ectotherms adapting to warmer temperatures ^31,32^. The fact that increasing the *T*_*pk*_s of certain traits relative to others is predicted to increase population fitness is also important from the perspective of constraints on evolution of thermal fitness. For example, when populations are confronted with long-term climatic warming (across generations), we expect the *T*_*pk*_s of development rate and maximum fecundity to shift first, and set the upper limit on how fast the local population can respond to temperature change. To test the theoretical prediction that the trait TPCs underlying thermal fitness have evolved in a hierarchical way, we performed an exhaustive data synthesis covering 61 diverse arthropod species (Fig. 3A; Methods). Although this empirical data synthesis largely exhausts the data available in the literature, this relatively small number of species highlights the relative paucity of data on arthropod thermal traits. Nevertheless, our synthesis provides TPC data for an unprecedented diversity of arthropod lineages, and allows us to test the generality of our results.

**Figure 3:**
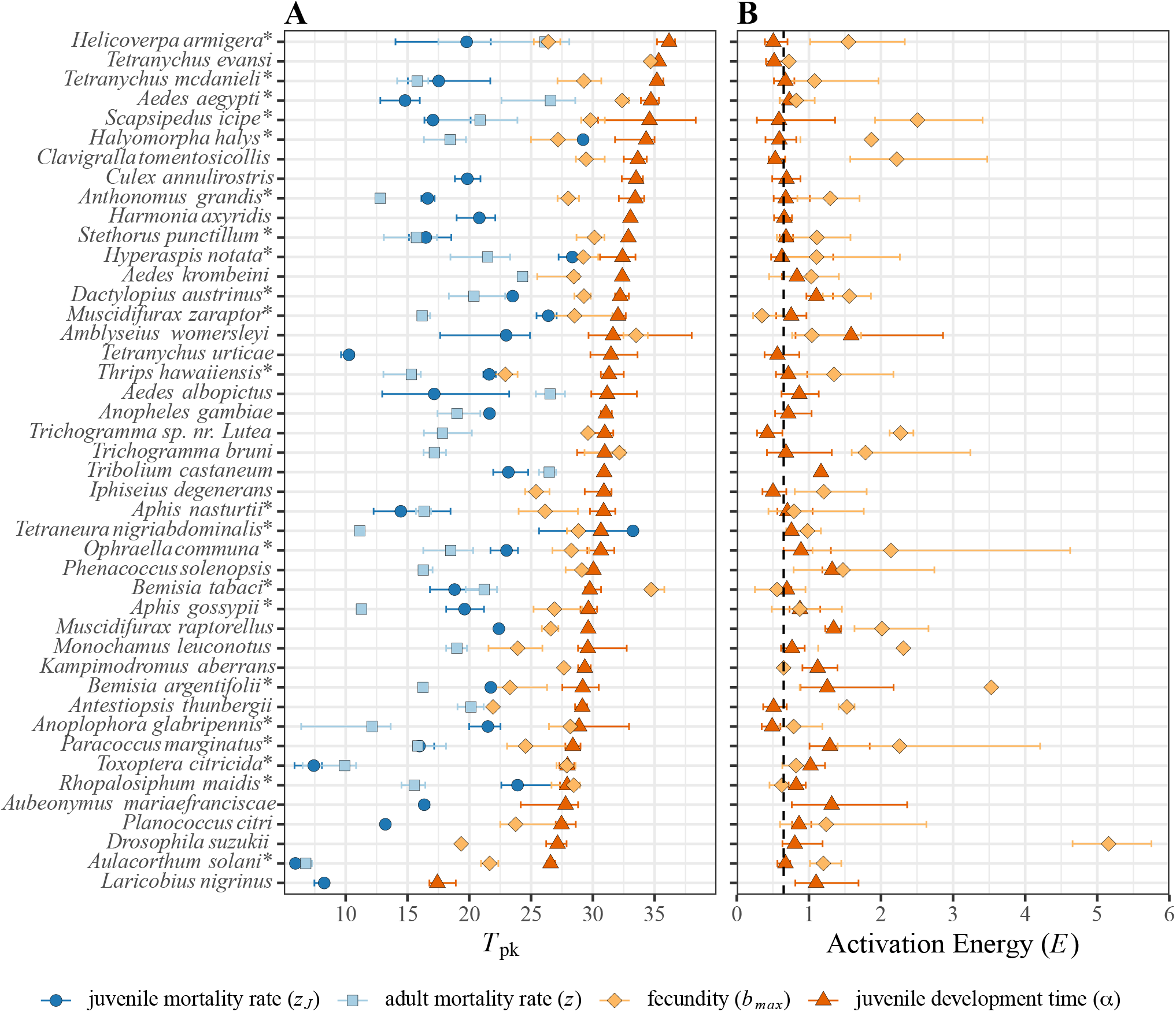
Empirical patterns in arthropod trait peak temperatures (*T*_*pk*_s) **(A) and thermal sensitivities (***E***s) (B)** (with 95% confidence intervals obtained by bootstrapping). Species for whom the full complement of trait data necessary for evaluating the *r*_*m*_ model (Methods, Equation (2)) are denoted with asterisks (*n*=22 out of 44 featured here). Species that did not have data for *α* (*n*=10) or only had *α* (*n*=7) are included in Supplementary Fig. 1.

We first tested for the predicted existence of a hierarchical ordering of within-species trait *T*_*pk*_s. We found that the empirical patterns are remarkably consistent with the prediction that taxa evolve to optimize their fitness by maximizing the *T*_*pk*_ of traits in a specific order (Fig. 2). Firstly, development rate almost always exhibits a higher *T*_*pk*_ than peak fecundity, adult mortality rate, or juvenile mortality rate (in all but three species; Fig. 3A). Secondly, in the 22 species for which we were able to find data for all four traits, the ordering of *T*_*pk*_s in 55% is exactly as predicted 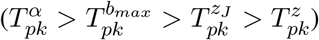. The probability of observing this ordering of trait *T*_*pk*_s by random chance is negligible. The match to our theoretical predictions is even stronger if we ignore the data on the trait under weakest selection (adult mortality rate), in which case 68% of the 44 species with data on all three of the more strongly-selected traits—development, peak fecundity and juvenile mortality rate—, show the expected ordering of *T*_*pk*_’s. Also, as expected from ecological metabolic theory ^10,14,23^, the thermal sensitivity parameters (*E*) of trait TPCs is relatively constrained across species, emphasising the primacy of evolution of trait *T*_*pk*_s relative to thermal sensitivity in driving thermal adaptation in arthropods (Fig. 3B).

To investigate the long-term evolution of the four trait *T*_*pk*_s across species, we then performed a phylogenetic analysis. The results again support the predicted hierarchy of selection on *T*_*pk*_s (Fig. 4A). Nevertheless, a given trait’s *T*_*pk*_ values for a closely related species pair were generally more similar than those for randomly selected pairs (high phylogenetic heritability in Fig. 4B). Thus, adaptive shifts in a trait’s *T*_*pk*_ within lineages have not overcome differences across distantly-related species even through long-term evolution. These differences may be ascribed to differences in metabolic (i.e., genomic) architecture between arthropod lineages ^33^. Phylogenetic heritability was the weakest for 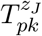, suggesting a more pronounced role of plasticity in the shift in its *T*_*pk*_ compared other traits, perhaps because of stage-specific mortality patterns (note that *z*_*J*_ is mortality rate across juvenile lifestages). We also found evidence that 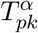has evolved more slowly than 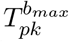(Fig. 4C), consistent with the predicted stronger (stabilising) selection on 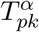 relative to the *T*_*pk*_s of the other traits (Fig. 2).

**Figure 4:**
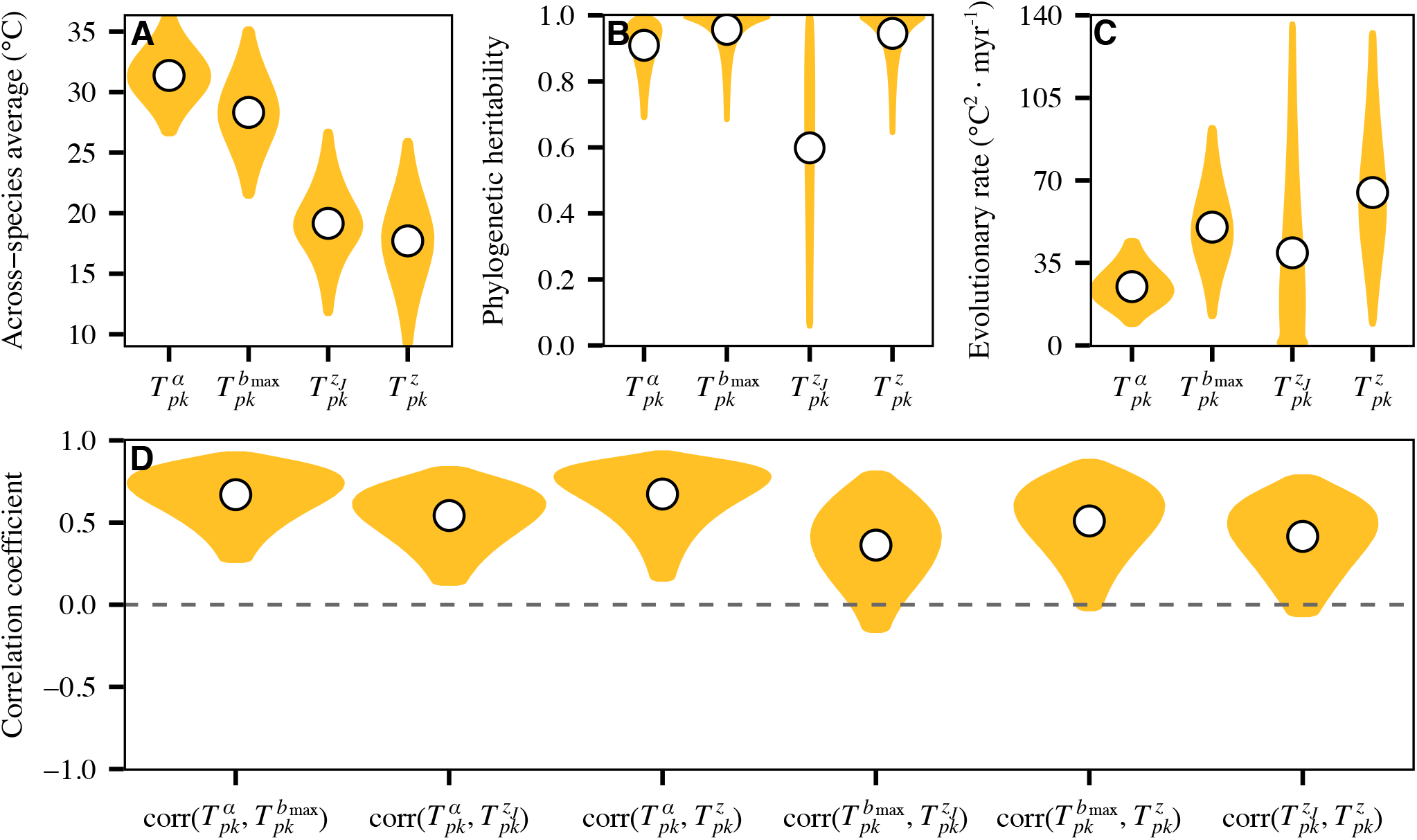
Macroevolutionary patterns of trait *T*_*pk*_s across arthropods. **A:** The average *T*_*pk*_ values across species after accounting for evolutionary relationships. **B:** The fraction of variance in *T*_*pk*_ values that is explained by the phylogeny assuming a Brownian Motion model of trait evolution. The remaining variance arises from phylogenetically-independent sources (e.g., plasticity). **C:** The median rates of evolution (under Brownian Motion) per million years. **D:** Median phylogenetic correlations between trait *T*_*pk*_ pairs. Gold areas in all plots represent the 95% Highest Posterior Density intervals (Methods).

### Physiological mismatches constrain population fitness

We next focused our analysis on the role of potential trade-offs between the *T*_*pk*_s of life history traits in constraining the optimization of thermal fitness. Our theory predicts that at a given temperature, the simultaneous maximisation of multiple trait *T*_*pk*_s should increase thermal fitness exponentially (Fig. 5A). That is, thermal fitness is exponentially higher if the *T*_*pk*_s of all traits increase in concert in a warmer environment. For example, if the development rate *T*_*pk*_ increases, the emergent thermal fitness will be higher if the *T*_*pk*_s of all other traits increased with it than if they did not. This result, along with our selection gradient analysis (Fig. 2), predicts that selection should favor not just a maximization of 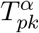 relative to the *T*_*pk*_s of other traits, but also a minimization of the differences (i.e., “physiological mismatches”) among the trait *T*_*pk*_s. However, the constraints of a fixed energy budget would limit such evolutionary optimisation, imposing trade-offs. For example, at a given temperature, available energy can be allocated to, say, either maximize development rate or fecundity rate, but not both ^34,35^.

**Figure 5:**
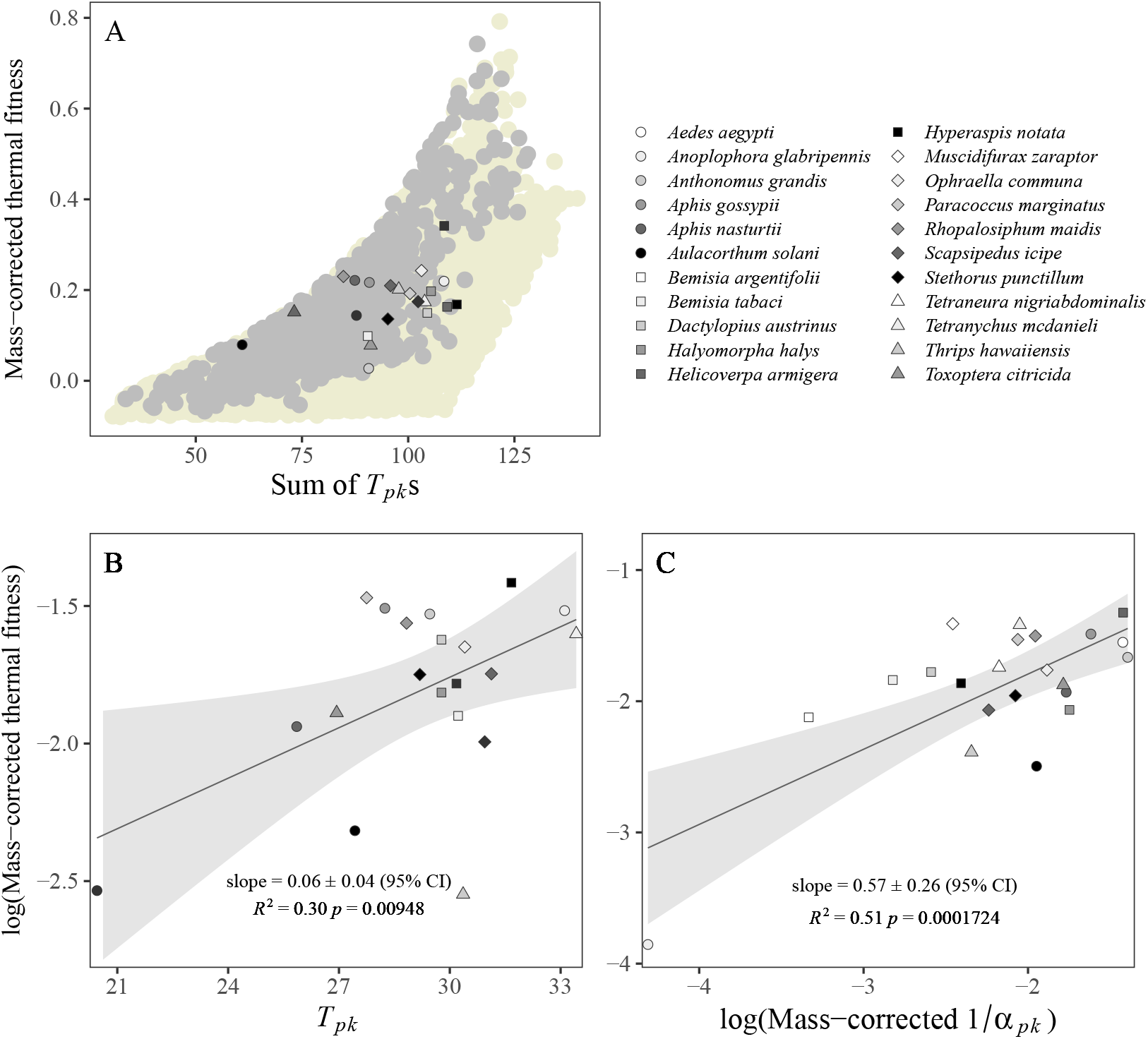
Physiological mismatches predict patterns of thermal fitness in extant arthopod taxa. **A:** Thermal fitness (*r*_*opt*_) increases with the sum of trait *T*_*pk*_s, which quantifies the simultaneous maximisation (optimization) of trait TPCs. The pale yellow dots represent simulated thermal fitness obtained by randomly varying the optimal temperatures (*T*_*pk*_s) of all traits 5000 times by drawing them from a uniform distribution between of 10 *−*35^*°*^C (Methods) without constraining the ordering of the trait *T*_*pk*_s. Among these, the subset of light grey dots satisfy the predicted hierarchy of trait selection strengths (are closer to an optimal strategy of 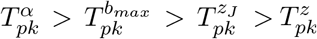) (Fig. 2). The thermal fitness values for 22 species (Fig. 3) with sufficient trait data are overlaid on these theoretically-predicted strategies. **B & C**: Thermal fitness increases with optimal temperature across a diverse group of arthropod taxa, i.e., a hotter-is-better pattern. **B:** Mass-corrected optimal *r*_*m*_ (*r*_*opt*_/*M* ^*−*0.1^) predicted by Equation (2) increases with its *T*_*opt*_. **C:** Consistent with our theoretical prediction, mass-corrected optimal thermal fitness (*r*_*opt*_/*M* ^*−*0.1^) increases linearly (in log-scale) with mass-corrected development rate *T*_*pk*_s ((1/*α*_*pk*_)/*M* ^*−*0.27^).

Indeed, our empirical data analyses show that there are substantial physiological mismatches in traits across diverse extant arthropod taxa (Fig. 3A). For example, the four species with the highest *T*_*pk*_s for development rate (*H. armigera, T. mcdanieli, A. aegypti*, and *S. icipe*) have as much as a *∼*20^*°*^C mismatch with the *T*_*pk*_s of juvenile mortality and adult mortality rates (which lie at the other end of the selection strength hierarchy; Fig. 2). This pattern suggests that the maximisation of performance of multiple traits at the same temperature is somehow constrained.

Our phylogenetic analyses also revealed significant correlated evolution between 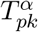 and the three other trait *T*_*pk*_s (Fig. 4D). This is in line with the theoretical prediction that multiple trait *T*_*pk*_s should move in concert (Fig. 5), but also reflect the fact that despite differences in genomic architecture, these four life-history traits likely share core metabolic pathways ^33,36^. Furthermore, that all the correlations are somewhere in the order of 0.5, is also in line with trade-offs limiting the degree of *T*_*pk*_ convergence across traits.

It is worth noting that these patterns of correlated evolution of trait *T*_*pk*_ are counter to the temperature-size rule ^37^, which states that body size decreases with temperature due to accelerated development. Based on the this rule we would instead predict a weak positive or even negative correlation between 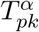 and 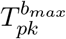. This is because of a life history trade-off: on the one hand, a larger body size increases fecundity and fitness, while on the other, growing for a longer duration increases realised mortality across juvenile stages and decreases the number of generations completed within a year or season, ultimately decreasing fitness. Indeed, a growing number of studies have found significant selection on larval development rate to reduce juvenile mortality and generation time, counterbalancing the positive fitness effect of increasing body size on fecundity ^38,39,40^. This apparent contradiction between the prediction of the size-temperature rule and the macroevolutionary patterns we report here can be reconciled by the fact that the size-temperature rule operates at shorter timescales, affording phenotypic plasticity to populations facing sudden temperature changes. Indeed, future climate change will likely entail not just directional warming but also increases in frequency and magnitude of thermal fluctuations (extreme events). Further work is needed to determine the relative role of phenotypic plasticity versus rapid evolution to build a more complete picture of the ability of arthropod populations to persist under future climate change ^17,18,41^. In that context, it is also worth noting that the optimal strategy for a population in fluctuating thermal environments is to evolve towards a relatively lower *T*_*opt*_ (and thus thermal fitness) in order to reach an adequate thermal safety margin ^41,42,43^. In that respect, extending our theoretical framework and thermal fitness calculations to account for fluctuating environments would likely yield important further insights into the constraints imposed by the evolution of trait-specific TPCs (Fig. 1, 2) on the adaptive potential of arthropod populations.

Finally, to test whether arthropod traits have nevertheless evolved to achieve maximal fitness within the constraints imposed by trade-offs, we overlaid the estimated thermal fitness from the data synthesis on to the theoreticallypredicted optimal region. We found that real thermal fitness values of diverse arthropod taxa do indeed lie within our theoretically-predicted ranges (Fig. 5A). Furthermore, this thermal life history optimization operates under fundamental thermodynamic constraints, reflected in a global hotter-is-better pattern of thermal fitness (Fig. 5B; note that the 22 species’ *r*_*opts*_ are within the global hotter-is-better pattern). Additionally, as expected, thermal fitness and its hotter-is-better pattern is strongly predicted by the (mass-corrected) value of development rate (1*/α*) at its *T*_*pk*_ (Fig. 5C; also see Supplementary Results Section 1.2). We also find that as expected (and assumed by our theory) the underlying traits (and most crucially, development time, *α*) too follow a hotter-is-better pattern (Supplementary Results Section 1.1). We also confirmed that these empirical patterns across diverse taxa are the result of long-term adaptive thermal evolution by analysing data on the native thermal environments of these taxa (Supplementary Results Section 1.3).

## Conclusions

We have developed a relatively simple theoretical framework to make meaningful predictions about the temperature dependence of population fitness, adaptation potential and climatic vulnerability across diverse arthropods. This framework requires data on the temperature-dependencies of just five key measurable life history traits across diverse arthropod taxa. This framework also allows constraints imposed by trade-offs between traits’ performances, well known to limit fitness optimization in other contexts ^44^, to be considered in the context of thermal adaptation in a nuanced yet general manner. Our overall approach and framework could thus guide conservation and risk assessment efforts by looking across species to identify those groups most vulnerable to extinction and those most likely to expand their distributions in the face of climate change.

## Methods

### The model for temperature-dependent maximal population growth rate

Our model for *r*_*m*_ is based on the Euler-Lotka equation ^14,45,46^:

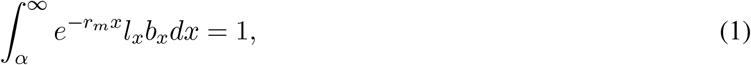

where *α* is the age at first reproduction (the time needed for development from egg to adult, or the juvenile to adult development time); *l*_*x*_ is age-specific survivorship (proportion of individuals that survive from birth to age *x*); and *b*_*x*_ the age-specific fecundity (number of offspring produced by an individual of age *x*). This equation gives the expected reproductive success of a newborn individual in a population growing at a rate *r*_*m*_, under the assumption that the population has reached a stable age distribution (i.e., the proportion of individuals in adult and juvenile life stages is constant). Using the simplest feasible mortality and fecundity models (for *l*_*x*_ and *b*_*x*_ respectively), we previously derived an approximation appropriate for the range of growth rates typically seen across arthropods ^15^ (Table 1):

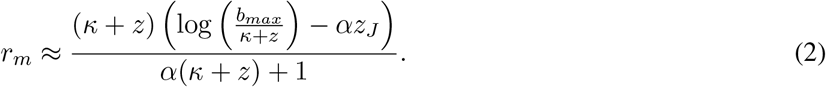

Substituting models of the thermal performance curves (TPCs) of the five life history traits (Supplementary Results, Section 1.5) into Equation (2) gives the temperature-dependence of *r*_*m*_. We model these traits TPCs using the modified Sharpe-Schoolfield equation ^23,27^ or its inverse (Main text, Fig. 1):

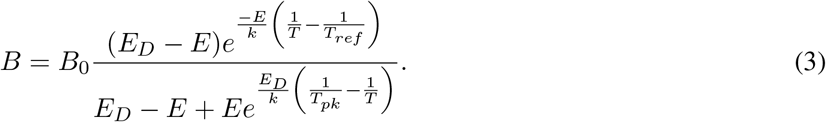

Here *B* is the value of the metabolic trait at a given temperature (*T*, in degrees K); *B*_0_ is a normalisation constant representing the value of the trait at some reference temperature (*T*_*ref*_); *E* is the apparent activation energy (initial thermal sensitivity), which determines how fast the curve rises up as the temperature approaches the peak temperature, *T*_*pk*_; and *E*_*D*_ is the deactivation energy, which determines how fast the trait declines after the peak.

The parameter *k* is the universal Boltzmann constant (8.617 · 10^*−*5^ eV *K*^*−*1^). The constant *B*_0_ also includes the effect of body size, and therefore the effect of stage-specific size differences, which we do not explicitly consider here theoretically, but will consider in the data we analyze (below; also see Discussion). Equation (3) has been used as a model for thermal performance of traits in numerous previous studies on arthropod population biology because it accurately captures the temperature dependence of a wide range of metabolically constrained life history traits ^10,29,47^. For mortality rates (*z* and *z*_*j*_), we use the inverse of Equation (3) because the thermal response of mortality rate tends to be U-shaped ^48,49,50^, and is well-fitted by it ^47,51^. Using a different unimodal function instead of the Sharpe-Schoolfield equation will not qualitatively change our results provided that it can encode a “hotter-is-better” constraint ^28^. Substituting the trait TPCs into Equation (2) yields the TPC of *r*_*m*_, from which we numerically calculate the *T*_*opt*_ (temperature at which the optimum of population growth occurs) and *r*_*opt*_ (the value of *r*_*m*_ at *T*_*opt*_). Note that we refer to the temperature of peak *r*_*m*_ as *T*_*opt*_ rather than *T*_*pk*_ as we do for traits, because the *T*_*pk*_s of those traits do not necessarily correspond to the thermal optimum of population fitness (optimal thermal fitness).

### Trait sensitivity analysis

To determine how much influence individual traits’ TPCs have on *r*_*m*_’s temperature-dependence, using the chain rule, we can write ^15,48^:

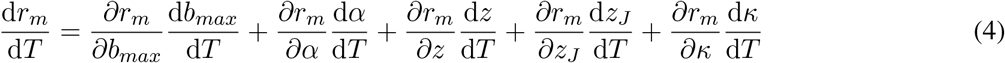

Each summed term in the right hand side of this equation quantifies the relative contribution of the TPC of a parameter to the temperature dependence of *r*_*m*_ (Main text, Fig. 1C). The trait TPCs for this calculation were parameterized as described above with and identical *T*_*pk*_ = 25^*°*^C across all traits. Here again, the choice of the TPC parameters, as long as all the *B*_0_s are in their appropriate scale, do not change the results of this trait-sensitivity analysis qualitatively.

### Quantifying trait-specific selection gradients

In order to quantify the impact of changes in *T*_*pk*_s of different traits on *r*_*m*_’s temperature-dependence (i.e., the thermal selection gradient on each trait), we calculated the shift in optimal maximum population growth rate (*r*_*opt*_), i.e., the value of *r*_*m*_ at *T*_*opt*_ with a unit change in each trait’s *T*_*pk*_. For this, re-evaluated Equation (2) by varying each trait’s *T*_*pk*_ in turn while holding all other trait TPCs constant at 25^*°*^C (approximately the median of all the values observed in the data; Main Text, Fig. 3), over a temperature range of 0–35^*°*^C (Main text, Fig. 2). That is, we allowed *T*_*pk*_ of each focal trait, in turn, to vary from -10 to +15 ^*°*^C relative to *T*_*pk*_ = 25^*°*^C, keeping all other trait TPCs (including their *T*_*p*_*k*s) fixed. This ensured that the *T*_*pk*_s of any given trait varied between 15^*°*^C – 40^*°*^C, which is approximately the range seen in the empirical data (see below; Main text, Fig. 3). We denote the difference of the focal trait’s *T*_*pk*_ from those of all others as 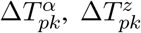, etc. The strength of the directional selection gradient for each trait was calculated as the second derivative 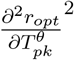 (where *θ* is one of the four traits) ^12,22,52,53^. The minimum of this quantity pinpoints the temperature at which that selection gradient starts to asymptote as serves as a prediction of the ordering of trait *T*_*pk*_s that optimises thermal fitness (Main text, Fig. 2). This method ignores trade-offs or covariances between traits, which would change the shape of the selection gradients (e.g., making them unimodal instead of asymptotic; see discussion in Main text, as well as the results of our phylogenetic analyses (Fig. 4)).

### Quantifying effects of physiological mismatches

Given that a higher *T*_*pk,α*_ increases fitness, and that all trait-specific selection gradients are monotonically increasing, closer the *T*_*pk*_s of the other traits are to *T*_*pk,α*_, the higher the *r*_*opt*_. We term the distance between any trait’s *T*_*pk*_ and *T*_*pk,α*_ a “physiological mismatch”. To quantify the effect of the overall level of physiological mismatch across all pairwise combinations of *T*_*pk,α*_ and each of the other three traits, we used the sum of all *T*_*pk*_s as a mismatch measure, which is a consistent measure because it increases monotonically with a decrease in mismatch. Then to calculate the effects of this overall physiological mismatch on thermal fitness, we quantified how *r*_*opt*_ changes with different potential life history strategies (in terms or ordering of the *T*_*pk*_s) it by randomly permuting their order. For this, all other TPC parameters were kept fixed (see next).

### TPC parameterisations

For the numerical calculation of *r*_*m*_’s temperature-dependence, trait sensitivity, selection gradient analyses and physiological mismatch analyses (above), we parameterized the traits’ TPC equations as follows. Firstly, because our theory focuses on the relative differences between the *T*_*pk*_s of traits, the exact values of *E* and *E*_*D*_ do not matter as long as they lie within empirically-reasonable values, so we fixed them across all traits to be *E* = 0.6 and *E*_*D*_ = 4. These are approximately the median values found in our empirical data (Main text Fig. 3, Supplementary Fig. 6). Varying these within the range seen in our empirical data do not change our results qualitatively. The parameter values for *B*_0_ were varied with type of trait, fixing them to be (approximately, rounded) the median values observed in our empirical data at a *T*_*ref*_ of 10^*°*^C: *B*_0,*α*_ = 25, 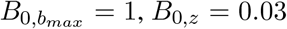, and 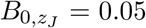 (Supplementary figures 7-10). In the absence of data on the rate of loss of fertility (*κ*) or its TPC, we evaluated the model’s predictions for a range of values of this parameter (Supplementary Results Section 1.7). While the values for all these parameters in the real data vary considerably around their respective medians, these specific ones used throughout the manuscript’s theory figures suffice to generate qualitatively-robust theoretical predictions.

### Data synthesis

We performed an extensive literature search to collate existing data on the TPCs of individual traits across arthropod taxa. We searched for publications up to July 2022 using Google Scholar’s advanced search, using Boolean operators (e.g., life history AND pest OR vector AND temper*), without language restrictions. Additional searches were also made by including species’ names in the search string to improve the detection of publications on underrepresented groups. For some species, multiple data sets were available so we only included the study that provided the most complete data (i.e., the highest number of traits measured). Raw data and references are available in Appendix 1. We excluded all field studies, and also lab studies where traits had been measured over narrow (*<* 10^*°*^C) temperature ranges (that prevented reliable TPC model fitting; below). We also excluded species from our analysis of physiological mismatches if TPC data for at least two traits were not available for them. Values of relevant traits across temperatures at were extracted from the text or tables, or were read from the figures using WebPlot-Digitizer ^54^. All data were converted to consistent measurement units. If juvenile mortality rate (*z*_*J*_, individual^*−*1^ day^*−*1^) was not provided (most of the original studies), we divided juvenile survival proportion by development time. Similarly, if adult mortality rate (*z*) was not directly reported, we calculated it as the reciprocal of (typically, female) longevity (1/days). When fecundity rate was not directly reported we divided lifetime reproductive output by longevity to obtain fecundity rate (*b*_*max*_; eggs / (individual*×* day)^*−*1^).

Finally, each species- and trait-specific thermal response dataset was fitted to the appropriate TPC equation (Equation (3) or its inverse; Fig. 1A & B) using NLLS using the `rTPC` pipeline ^30,55^. Bootstrapping (residual resampling) was used to calculate 95% prediction bounds for each TPC, which also yielded the confidence intervals around each *T*_*pk*_ and peak trait value (*B*_*pk*_) estimate. These trait value estimates were body mass-corrected to adjust for how the temperature-dependence of *r*_*m*_ emerges from the temperature-dependencies of its underlying traits, which, in turn, are dependent on the relationships between body size and metabolic rate in individual organisms ^46^. If fresh mass (mg, *M* in Main text Fig.4; Supplementary figures 1 & 2) for a particular species was not provided in the original study, we used mass estimates from other studies on that or a closely related species.

### Phylogenetic analyses

#### Phylogeny construction

To examine the macroevolutionary patterns of the *T*_*pk*_s of the four main traits of this study (*α, b*_*max*_, *z*_*J*_, and *z*), we first extracted the phylogenetic topology of all the species in our dataset from the Open Tree of Life ^56^ (OTL; v.13.4) using the `rotl` R package ^57^ (v.3.0.12). Given that the OTL topology (Supplementary Fig. 16A) included a few polytomies, we also collected publicly available nucleotide sequences (where available) of: i) the 5’ region of the cytochrome c oxidase subunit I gene (COI-5P); ii) the small subunit rRNA gene (SSU); and iii) the large subunit rRNA gene (LSU). COI-5P sequences were obtained from the Barcode of Life Data System database ^58^, whereas SSU and LSU sequences were extracted from the SILVA database ^59^ (Supplementary Table 1).

Next we aligned the sequences with MAFFT ^60^ (v.7.490) using the G-INS-i algorithm for COI-5P sequences, and the X-INS-i algorithm for SSU and LSU sequences. We specifically chose the latter algorithm for the two rRNA genes as it can take the secondary structure of RNA into consideration while estimating the alignment ^61^. We then removed phylogenetically uninformative sites using the Noisy tool ^62^, and merged the alignments of the three genes into a single concatenated alignment. For each gene, we identified the optimal model of nucleotide substitution with ModelTest-NG ^63,64^ (v.0.2.0), according to the small sample size-corrected Akaike Information criterion ^65^.

For phylogenetic topology inference based on the concatenated alignment, we employed the RAxML-NG tool ^66^ (v.1.1.0). We constrained the topology search based on the OTL tree, which allowed us to incorporate further phylogenetic information from previously published studies. Nevertheless, because we could not obtain molecular sequences for all species (Supplementary Table 1), some of the polytomies of the OTL tree could not be objectively resolved. To account for this uncertainty in downstream analyses, we performed 100 topology searches, each of which started from 100 random and 100 maximum parsimony trees. This process yielded a set of 100 trees, in which 18 alternative topologies were represented (Supplementary Fig. 16B).

To time-calibrate each of the 100 trees, we first queried the Timetree database ^67^ to obtain reliable age information (based on at least five studies) for as many nodes as possible. We then applied the “congruification” approach ^68^ implemented in the `geiger` R package ^69^ (v.2.0.10). In other words, we transferred known node ages (and their uncertainty intervals) from the reference phylogeny (TimeTree) to each of the 100 target phylogenies. This information was then used by the treePL tool ^70^ to estimate ages for all tree nodes based on penalized likelihood. Supplementary Fig. 16B shows the final set of the 100 alternative time-calibrated trees.

#### Investigation of the macroevolutionary patterns of *T*_*pk*_s

To quantitatively characterise the evolution of the *T*_*pk*_s of the four traits in this study, we fitted a Bayesian phylogenetic multi-response regression model with the `MCMCglmm` R package ^71^ (v.2.33). In particular, this model had all four *T*_*pk*_s as separate response variables and one intercept per response. This allowed us to simultaneously estimate both the variances and covariances among the four *T*_*pk*_s and, through this, to detect any systematic correlations. Furthermore, we specified a phylogenetic random effect on the intercepts by integrating the phylogenetic variance-covariance matrix into the model. By doing so, we partitioned the variance-covariance matrix of *T*_*pk*_s into a phylogenetically heritable component and a residual component. The latter would reflect phylogenetically independent sources of variation in *T*_*pk*_ values (e.g., plasticity, experimental noise).

Given that we had information on the uncertainty of each *T*_*pk*_ estimate from bootstrapping, we directly incorporated this into the model. Missing *T*_*pk*_ values for one or more traits per species were modelled as “Missing At Random” ^72,73^. This approach (not to be confused with “Missing Completely At Random”) allows missing values in a response variable to be estimated (with some degree of uncertainty) from other covarying variables and from the phylogeny, provided that missingness is not systematically driven by a variable that is not included in the model (e.g., body size, habitat). Lastly, we specified uninformative priors, namely the default normal prior for the fixed effects, a Cauchy prior for the random effects covariance matrix, and an inverse Gamma prior for the residual covariance matrix.

We fitted this model 100 times, each time with a different phylogenetic tree from our set, and with three independent chains per tree. We set the chain length to two hundred million generations and recorded posterior samples every five thousand generations, except for the first 10% of each chain (i.e., twenty million generations) which we discarded as burn-in. We then verified that the three chains per tree had statistically indistinguishable posterior distributions and had sufficiently explored the parameter space. For these, we ensured that the potential scale reduction factor value of each parameter was smaller than 1.1, and that its effective sample size was at least equal to 1,000.

Finally, we combined the posterior samples from the three chains of all 100 runs. From these, we first extracted the intercept and the phylogenetically heritable variance of each *T*_*pk*_. The former represents the across-species mean whereas the latter corresponds to the evolutionary rate per million years. We additionally calculated the pairwise correlations between *T*_*pk*_s and their phylogenetic heritabilities, that is, the ratio of the phylogenetically heritable variance to the sum of phylogenetically heritable and residual variances. We summarised these parameters by calculating the median value and the 95% Highest Posterior Density interval.

## Data and code availability

All data and code for reproducing the study’s analyses can be found at https://github.com/PawarLab/TraitMismatchPaper-main.git. Our global dataset on arthropods is also available as an Auxiliary Supplementary file (Appendix 1).

## Author contributions

SP and LC conceived the study. SP developed the theory with support from TS and LJ. PH collated and analysed the data with support from AC, MN, and MS. DGK performed the phylogenetic analyses. All authors contributed to writing and revising the manuscript.

## References

1. Bar-On, Y. M., Phillips, R. & Milo, R. The biomass distribution on Earth. Proc. Natl. Acad. Sci. U. S. A. 115, 6506–6511 (2018).

2. Wagner, D. L., Grames, E. M., Forister, M. L., Berenbaum, M. R. & Stopak, D. Insect decline in the Anthropocene: Death by a thousand cuts. Proc. Natl. Acad. Sci. U. S. A. 118, 1–10 (2021).

3. Van Klink, R. et al. Meta-analysis reveals declines in terrestrial but increases in freshwater insect abundances. Science 368, 417–420 (2020).

4. Crossley, M. S. et al. No net insect abundance and diversity declines across US Long Term Ecological Research sites. Nat. Ecol. Evol 4, 1368–1376 (2020).

5. Marta, S., Brunetti, M., Manenti, R., Provenzale, A. & Ficetola, G. F. Climate and land-use changes drive biodiversity turnover in arthropod assemblages over 150 years. Nat. Ecol. Evol 5, 1291–1300 (2021).

6. Harvey, J. A. et al. Scientists’ warning on climate change and insects. Ecological monographs 93, e1553 (2023).

7. Heath, J. E., Hanegan, J. L., Wilkin, P. J. & Heath, M. S. Adaptation of the thermal responses of insects. Integr. Comp. Biol. 11, 147–158 (1971).

8. Jensen, A., Alemu, T., Alemneh, T., Pertoldi, C. & Bahrndorff, S. Thermal acclimation and adaptation across populations in a broadly distributed soil arthropod. Funct. Ecol. 33, 833–845 (2019).

9. Hoffmann, A. A. & Sgrò, C. M. Climate change and evolutionary adaptation. Nature 470, 479–85 (2011).

10. Dell, A. I., Pawar, S. & Savage, V. M. Systematic variation in the temperature dependence of physiological and ecological traits. Proc. Natl. Acad. Sci. U. S. A. 108, 10591–10596 (2011).

11. Frazier, M., Huey, R. B. & Berrigan, D. Thermodynamics constrains the evolution of insect population growth rates:”warmer is better”. Am. Nat. 168, 512–520 (2006).

12. Angilletta Jr, M. J. Thermal adaptation: a theoretical and empirical synthesis (Oxford University Press, 2009).

13. Kingsolver, J. G. et al. Complex life cycles and the responses of insects to climate change. Integr. Comp. Biol. 51, 719–732 (2011).

14. Amarasekare, P. & Savage, V. A framework for elucidating the temperature dependence of fitness. Am. Nat. 179, 178–91 (2012).

15. Cator, L. J. et al. The role of vector trait variation in vector-borne disease dynamics. Front. Ecol. Evol. 8, 189 (2020).

16. Jørgensen, L. B., Ørsted, M., Malte, H., Wang, T. & Overgaard, J. Extreme escalation of heat failure rates in ectotherms with global warming. Nature 1–6 (2022).

17. Duffy, K., Gouhier, T. C. & Ganguly, A. R. Climate-mediated shifts in temperature fluctuations promote extinction risk. Nat. Clim. Change 1–21 (2022).

18. Weaving, H., Terblanche, J. S., Pottier, P. & English, S. Meta-analysis reveals weak but pervasive plasticity in insect thermal limits. Nat. Commun. 13, 5292 (2022).

19. Deutsch, C. A. et al. Impacts of climate warming on terrestrial ectotherms across latitude. Proc. Natl. Acad. Sci. U. S. A 105, 6668–6672 (2008).

20. Brass, D. P. et al. Phenotypic plasticity as a cause and consequence of population dynamics. Ecol. Lett. 24, 2406–2417 (2021).

21. Buckley, L. B. & Kingsolver, J. G. Evolution of thermal sensitivity in changing and variable climates. Annu. Rev. Ecol. Evol. Syst. 52, 563–586 (2021).

22. Kingsolver, J. G. The well-temperatured biologist. Am. Nat. 174, 755–768 (2009).

23. Kontopoulos, D.-G. et al. Phytoplankton thermal responses adapt in the absence of hard thermodynamic constraints. Evolution 74, 775–790 (2020).

24. Sinclair, B. J., Williams, C. M. & Terblanche, J. S. Variation in thermal performance among insect populations. Physiol. Biochem. Zool. 85, 594–606 (2012).

25. Maino, J. L., Kong, J. D., Hoffmann, A. A., Barton, M. G. & Kearney, M. R. Mechanistic models for predicting insect responses to climate change. Curr. Opin. Insect Sci. 17, 81–86 (2016).

26. Angilletta, M. J., Huey, R. B. & Frazier, M. R. Thermodynamic effects on organismal performance: Is hotter better? Physiol. Biochem. Zool. 83, 197–206 (2010).

27. Schoolfield, R., Sharpe, P. & Magnuson, C. Non-linear regression of biological temperature-dependent rate models based on absolute reaction-rate theory. J. Theor. Biol. 88, 719–731 (1981).

28. Asbury, D. A. & Angilletta, M. J. Thermodynamic effects on the evolution of performance curves. Am. Nat. 176, E40–E49 (2010).

29. Huxley, P. J., Murray, K. A., Pawar, S. & Cator, L. J. The effect of resource limitation on the temperature dependence of mosquito population fitness. Proc. R. Soc. B 288, rspb.2020.3217 (2021).

30. Huxley, P. J., Murray, K. A., Pawar, S. & Cator, L. J. Competition and resource depletion shape the thermal response of population fitness in Aedes aegypti. Commun. Biol 5, 1–11 (2022).

31. Huey, R. B. & Berrigan, D. Temperature, demography, and ectotherm fitness. Am. Nat. 158, 204–210 (2001).

32. Trudgill, D. L. Why do tropical poikilothermic organisms tend to have higher threshold temperatures for development than temperate ones? Funct. Ecol. 9, 136–137 (1995).

33. Alfsnes, K., Leinaas, H. P. & Hessen, D. O. Genome size in arthropods; different roles of phylogeny, habitat and life history in insects and crustaceans. Ecology and Evolution 7, 5939–5947 (2017).

34. Tüzün, N. & Stoks, R. A fast pace-of-life is traded off against a high thermal performance. Proc. R. Soc. B (2022).

35. Birch, L. C. The intrinsic rate of natural increase of an insect population. J. Anim. Ecol. 17, 15 (1948).

36. Thomas, G. W. C. et al. Gene content evolution in the arthropods. Genome Biology 21, 15 (2020).

37. Atkinson, D. Temperature and organism size—a biological law for ectotherms? Advances in Ecological Research 25, 1–58 (1994).

38. Eck, D. J., Shaw, R. G., Geyer, C. J. & Kingsolver, J. G. An integrated analysis of phenotypic selection on insect body size and development time. Evolution 69, 2525–2532 (2015).

39. Huang, X.-L., Xiao, L., He, H.-M. & Xue, F.-S. Effect of rearing conditions on the correlation between larval development time and pupal weight of the rice stem borer, Chilo suppressalis. Ecology and Evolution 8, 12694–12701 (2018).

40. Chirgwin, E. & Monro, K. Correlational selection on size and development time is inconsistent across early life stages. Evolutionary Ecology 34, 681–691 (2020).

41. Dowd, W. W., King, F. A. & Denny, M. W. Thermal variation, thermal extremes and the physiological performance of individuals. J. Exp. Biol. 218, 1956–1967 (2015).

42. Martin, T. L. & Huey, R. B. Why “suboptimal” is optimal: Jensen’s inequality and ectotherm thermal preferences. Am. Nat. 171, E102–E118 (2008).

43. Bernhardt, J. R., Sunday, J. M., Thompson, P. L. & O’Connor, M. I. Nonlinear averaging of thermal experience predicts population growth rates in a thermally variable environment. Proc. R. Soc. B Biol. Sci. 285, 20181076 (2018).

44. Stearns, S. C. Trade-offs in life-history evolution. Funct. Ecol. 3, 259–268 (1989).

45. Charnov, E. L. Life history invariants: some explorations of symmetry in evolutionary ecology (Oxford University Press, Oxford England, 1993).

46. Savage, V. M. et al. Effects of body size and temperature on population growth. Am. Nat. 163, 429–41 (2004).

47. Molnár, P. K. P., Kutz, S. J. S., Hoar, B. M. B. & Dobson, A. P. A. A. P. Metabolic approaches to understanding climate change impacts on seasonal host-macroparasite dynamics. Ecol. Lett. 16, 9–21 (2013).

48. Mordecai, E. et al. Optimal temperature for malaria transmission is dramatically lower than previously predicted. Ecol. Lett. 16, 22–30 (2013).

49. Amarasekare, P. & Sifuentes, R. Elucidating the temperature response of survivorship in insects. Funct. Ecol. 26, 959–968 (2012).

50. Lunde, T. M., Bayoh, M. N. & Lindtjørn, B. How malaria models relate temperature to malaria transmission. Parasit. Vectors 6 (2013).

51. Van der Have, T. A proximate model for thermal tolerance in ectotherms. Oikos 98, 141–155 (2002).

52. Caswell, H. Matrix population models (Sinauer Associates Associates, Inc., Massachusetts, Massachusetts, 1989).

53. Hamilton, W. D. The moulding of senescence by natural selection. J. Theor. Biol. 12, 12–45 (1966).

54. Rohatgi, A. Webplotdigitizer: Version 4.5 (2021). URL https://automeris.io/WebPlotDigitizer.

55. Padfield, D., O’Sullivan, H. & Pawar, S. rTPC and nls. multstart: a new pipeline to fit thermal performance curves in R. Methods in Ecology and Evolution 12, 1138–1143 (2021).

56. Hinchliff, C. E. et al. Synthesis of phylogeny and taxonomy into a comprehensive tree of life. Proceedings of the National Academy of Sciences 112, 12764–12769 (2015).

57. Michonneau, F., Brown, J. W. & Winter, D. J. rotl: an R package to interact with the Open Tree of Life data. Methods in Ecology and Evolution 7, 1476–1481 (2016).

58. Ratnasingham, S. & Hebert, P. D. N. BOLD: The Barcode of Life Data System (http://www.barcodinglife.org). Molecular Ecology Notes 7, p355–364 (2007).

59. Quast, C. et al. The SILVA ribosomal RNA gene database project: improved data processing and web-based tools. Nucleic Acids Research 41, D590–D596 (2012).

60. Katoh, K. & Standley, D. M. MAFFT multiple sequence alignment software version 7: improvements in performance and usability. Molecular Biology and Evolution 30, 772–780 (2013).

61. Katoh, K. & Toh, H. Improved accuracy of multiple ncRNA alignment by incorporating structural information into a MAFFT-based framework. BMC Bioinformatics 9, 1–13 (2008).

62. Dress, A. W. M. et al. Noisy: identification of problematic columns in multiple sequence alignments. Algorithms for Molecular Biology 3, 7 (2008).

63. Darriba, D. et al. ModelTest-NG: a new and scalable tool for the selection of DNA and protein evolutionary models. Molecular Biology and Evolution 37, 291–294 (2020).

64. Flouri, T. et al. The phylogenetic likelihood library. Systematic Biology 64, 356–362 (2015).

65. Sugiura, N. Further analysis of the data by Akaike’s information criterion and the finite corrections. Communications in Statistics - Theory and Methods 7, 13–26 (1978).

66. Kozlov, A. M., Darriba, D., Flouri, T., Morel, B. & Stamatakis, A. RAxML-NG: a fast, scalable and user-friendly tool for maximum likelihood phylogenetic inference. Bioinformatics 35, 4453–4455 (2019).

67. Kumar, S. et al. TimeTree 5: An Expanded Resource for Species Divergence Times. Molecular Biology and Evolution 39, msac174 (2022).

68. Eastman, J. M., Harmon, L. J. & Tank, D. C. Congruification: support for time scaling large phylogenetic trees. Methods in Ecology and Evolution 4, 688–691 (2013).

69. Pennell, M. W. et al. geiger v2.0: an expanded suite of methods for fitting macroevolutionary models to phylogenetic trees. Bioinformatics 30, 2216–2218 (2014).

70. Smith, S. A. & O’Meara, B. C. treePL: divergence time estimation using penalized likelihood for large phylogenies. Bioinformatics 28, 2689–2690 (2012).

71. Hadfield, J. D. MCMC Methods for Multi-Response Generalized Linear Mixed Models: The MCMCglmm R Package. Journal of Statistical Software 33, 1–22 (2010).

72. Hadfield, J. & Nakagawa, S. General quantitative genetic methods for comparative biology: phylogenies, taxonomies and multi-trait models for continuous and categorical characters. Journal of Evolutionary Biology 23, 494–508 (2010).

73. de Villemereuil, P. & Nakagawa, S. General quantitative genetic methods for comparative biology. In Modern Phylogenetic Comparative Methods and Their Application in Evolutionary Biology: Concepts and Practice (ed. Garamszegi, L. Z.), 287–303 (Springer, 2014).

