## Supplemental Materials for "Variation in temperature of peak trait performance constrains adaptation of arthropod populations to climatic warming"

### Supplementary Materials for
### Variation in temperature of peak trait performance will constrain adaptation of arthropod populations to climatic warming

Samraat Pawar *et al.*
Paul J. Huxley,; Lauren Cator,

May 15, 2023

### 1 Supplementary Results

#### 1.1 Trait-level "hotter-is-better" patterns

Across diverse levels of biological organisation, many biological rates (e.g., development, population growth) are expected to scale to the negative quarter-power with mass-specific metabolic rate ($M^{-0.25}$) but such scaling may not exist in arthropods [1]. Therefore, prior to testing for trait-level "hotter-is-better" patterns, we fitted OLS models in log-log scale to the $B_{pk}$ estimates (Main text Fig. 4 and SM Figs. 1 & 2) and the body mass (wet mass, mg) data for each species (Appendix 1). We also size-corrected optimal thermal fitness ($r_{m,opt}$) to account for size scaling in underlying traits (Main text Fig. 4 and SM Fig. 2).

Our theory assumes that a "hotter-is-better" pattern exists in the underlying traits. In particular, given its dominant effect, $\alpha$ should definitely exhibit this pattern. While data on this within-species are not currently available, we tested this assumption with data across species (Main Text Fig. 4B; SM Fig. 1). We observe a significant "hotter-is-better" pattern for development time ($\alpha$; Main text Fig. 4B) and maximum fecundity ($b_{max}$) (SM Fig. 1A and D), whereas the slopes for $z_J$ and $z$ are non significantly positive and negative, respectively. *Anoplophora glabripennis* was excluded from this analysis because it's $r_{opt}$ was extremely low ($r_{opt}$=0.01). With this species included, the slope for seen in main text Fig. 4C becomes $0.09 \pm 0.08$ (95% CI), $R^2$=0.21 and $p$=0.03.

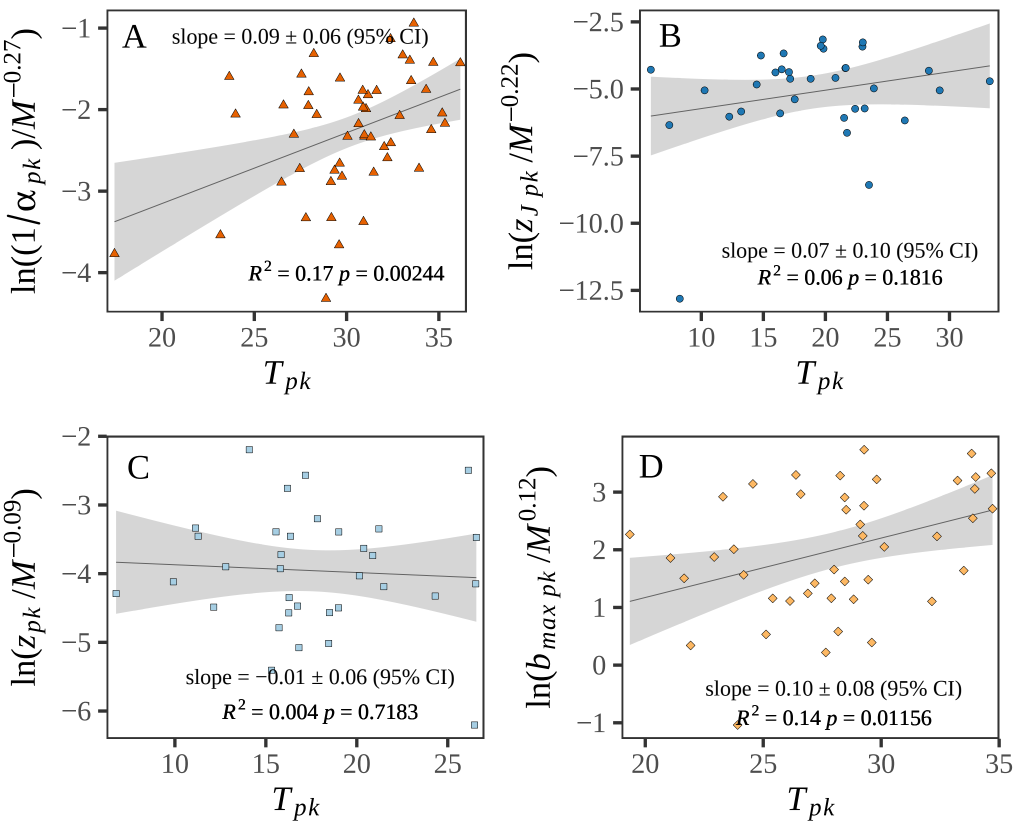

**Figure 1: Test of the "hotter-is-better" pattern across arthropod taxa**.The "hotter-is-better" pattern is significant in the body size-corrected (wet mass, mg) $1/\alpha$ (A) and $b_{max}$ (D) data, but at best weak for $z_J$ (B) and $z$ (C), suggesting relatively greater biochemical adaptation to overcome thermodynamic constraints in these two traits [2, 3] , insufficient data (note the narrow range of temperatures on x-axis for $b_{max}$ in particular) [4], or both. The lines are OLS regression (with 95% prediction bounds) fitted to log-transformed $B_{pk}$s of the trait plotted against the respective $T_{pk}$s.

#### 1.2 Correlation of thermal fitness with peak trait performance

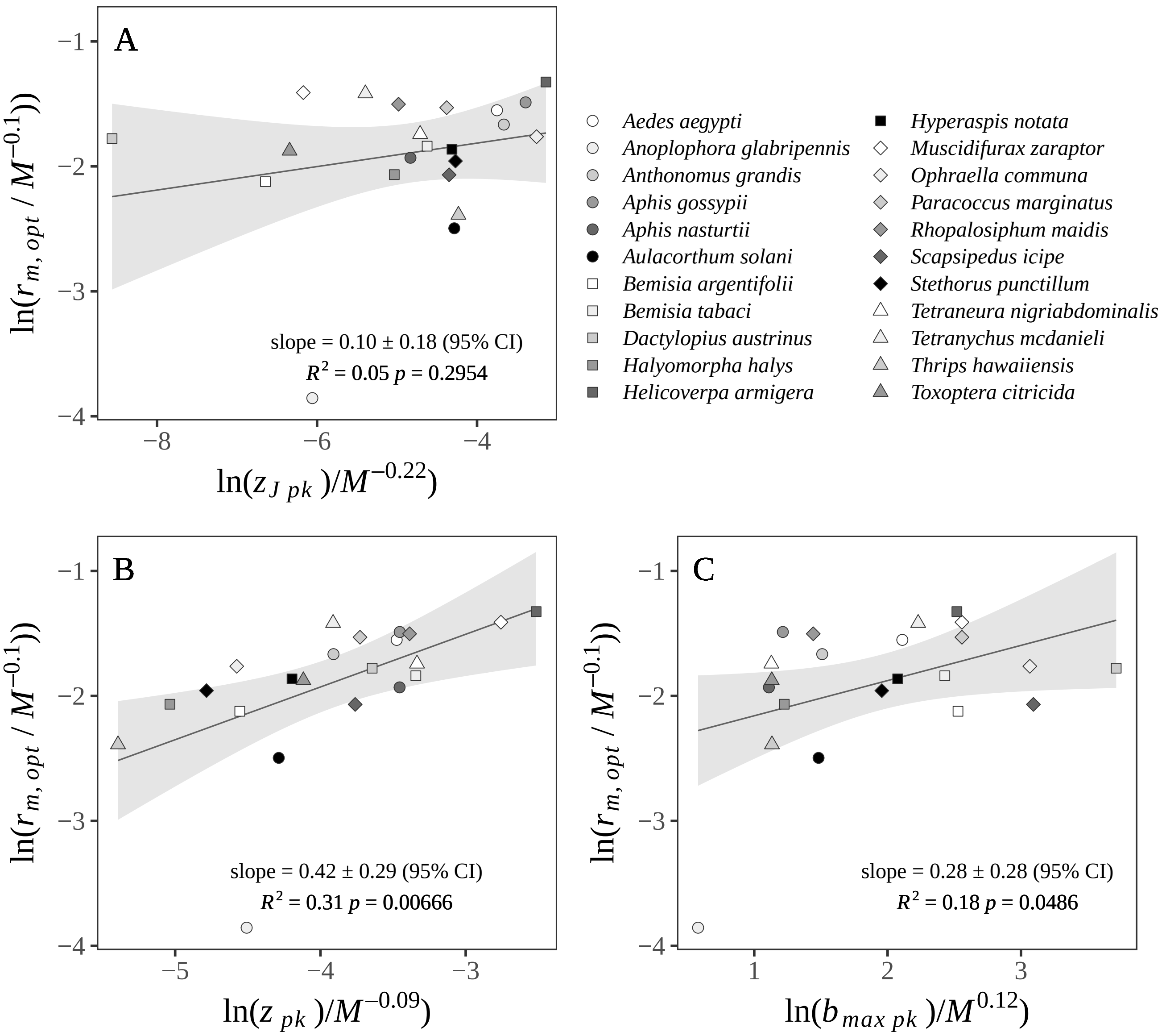

**Figure 2:** Relationship between $r_{m,opt}$ and peak A) juvenile mortality rate, B) adult mortality rate and C) fecundity rate. The lines are OLS regression (with 95% prediction bounds) fitted to log-transformed mass-corrected $r_{m,opt}$ plotted against the respective log-transformed mass-corrected trait $T_{pk}$s.

#### 1.3 Evidence of trait-level thermal adaptation

To test for trait-level thermal adaptation, we first analysed the relationships between trait $T_{pk}$s and latitudes (i.e., the geographical locations where each experimental species originated from). All traits' $T_{pk}$s declined with increasing latitude, which suggests that species, albeit weakly, are adapted to their local environments (SM Fig. 3). Because environmental variables other than temperature too vary with latitude we then also analysed the relationship between traits' $T_{pk}$s and rearing temperatures (i.e., the species' laboratory rearing temperatures; SM Fig. 4), and rearing temperatures and latitudes (SM Fig. 5). Traits' $T_{pk}$s increased significantly with rearing temperatures (SM Fig. 4) and decreased significantly as latitudes increased. These results suggest that species originating from warmer climates were reared under warmer temperatures in the lab. Together, these results indicate the existence of significant existing trait-level thermal adaption amongst the species used in the present study.

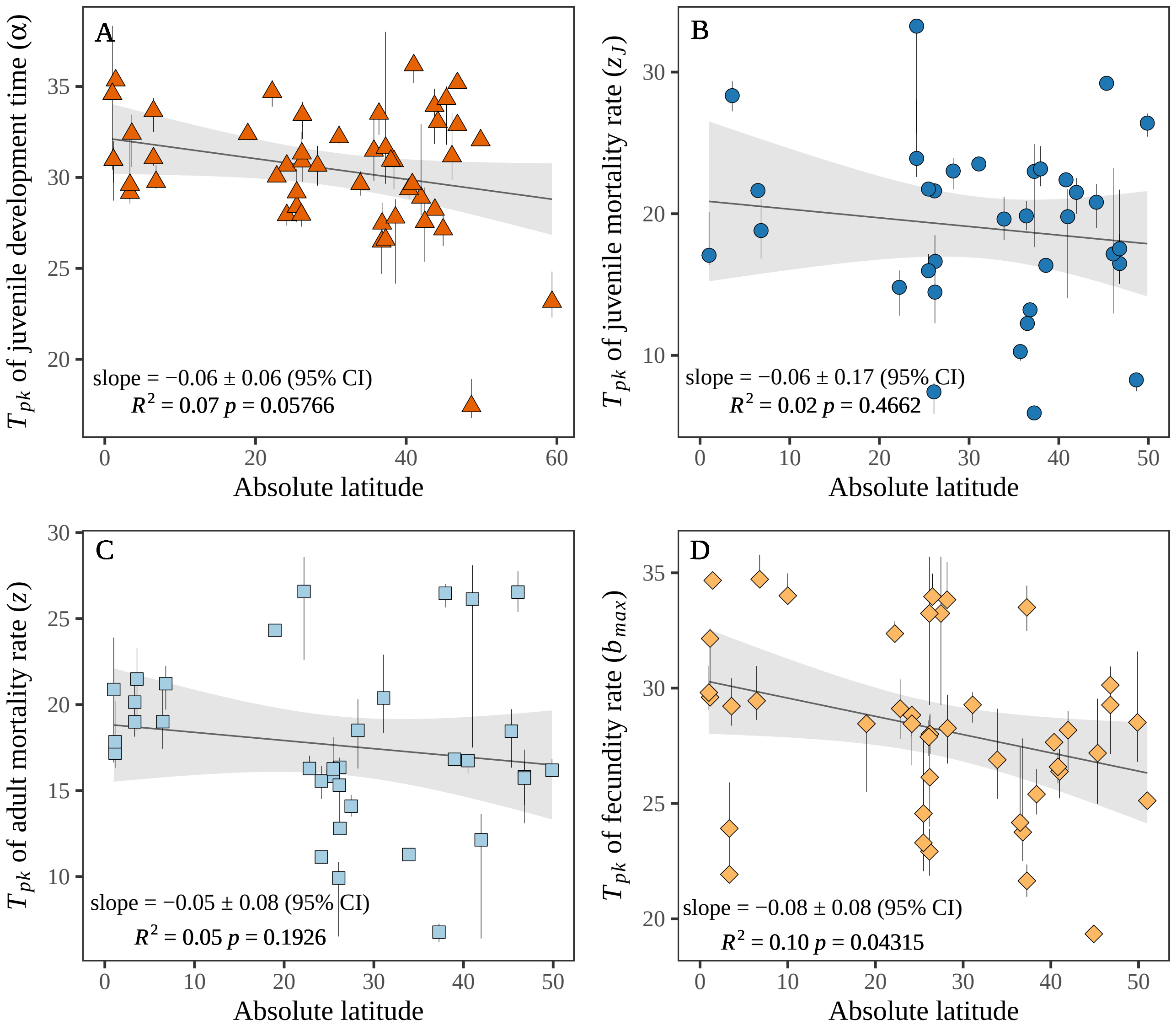

**Figure 3:** Relationship between latitude and $T_{pk}$s of A) juvenile development time, B) juvenile mortality rate, C) adult mortality rate and D) fecundity rate

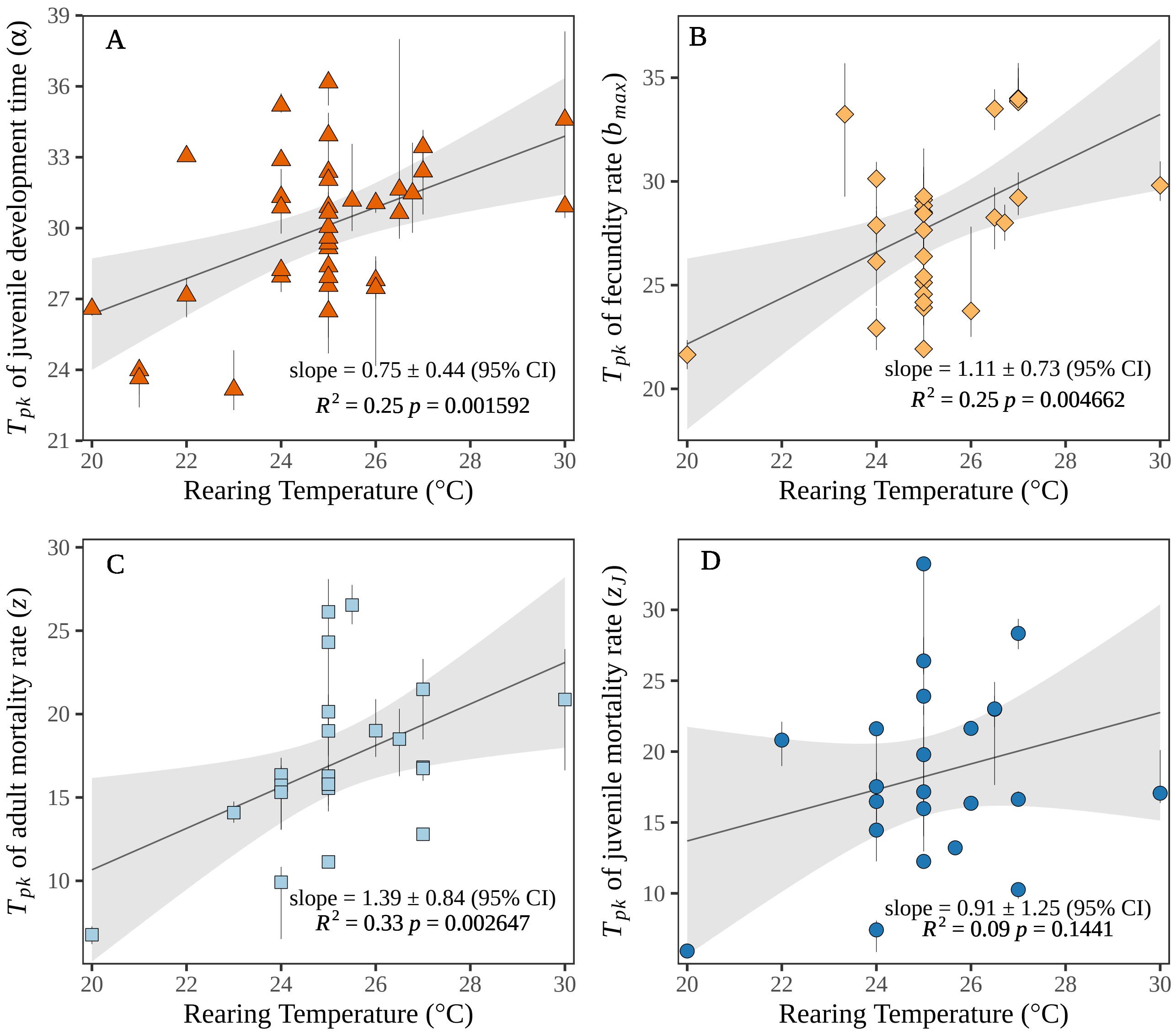

**Figure 4:** Relationship between rearing temperature and $T_{pk}$s of A) juvenile development time, B) juvenile mortality rate, C) adult mortality rate and D) fecundity rate

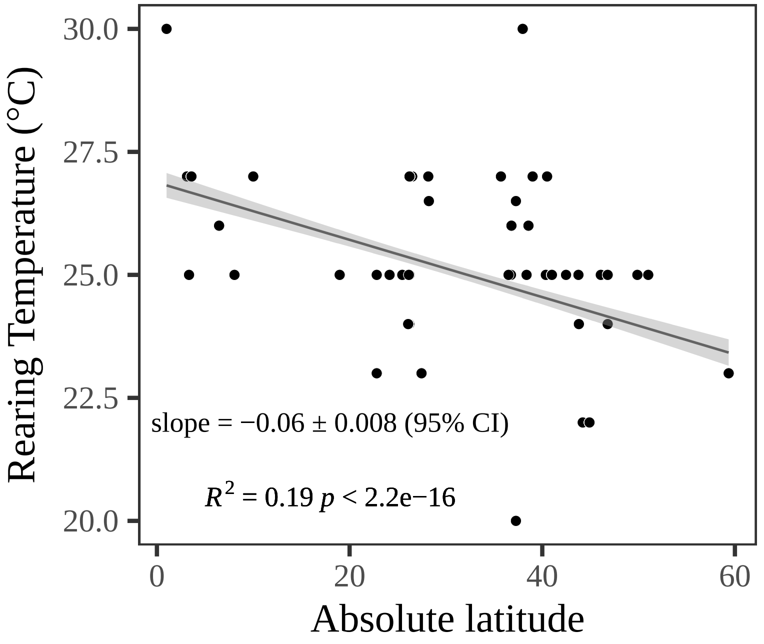

**Figure 5:** Relationship between rearing temperature and absolute latitude

#### 1.4 Distributions of trait-level thermal sensitivities

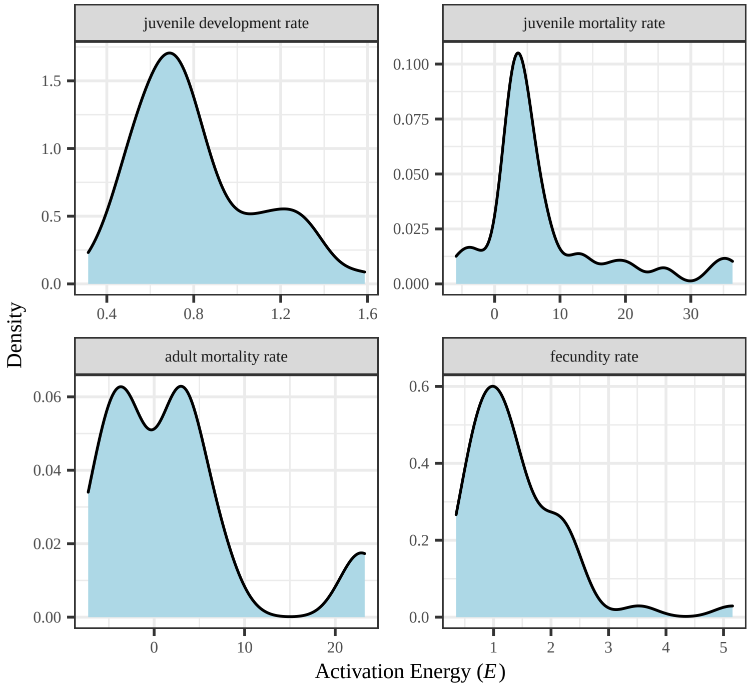

**Figure 6:** Distribution of estimated activation energy values for all fitted traits and species.

#### 1.5 Trait-level thermal performance curves

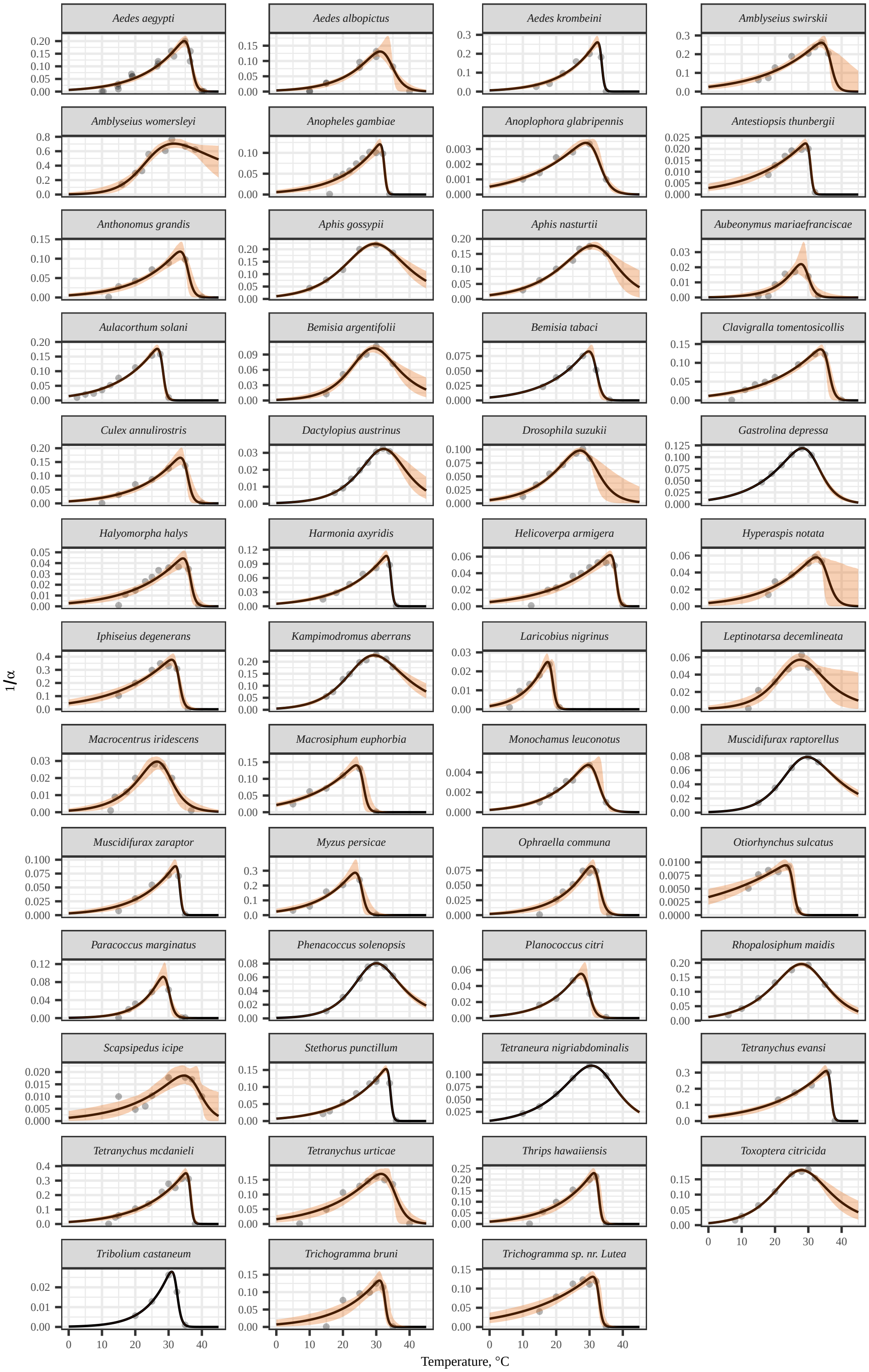

**Figure 7:** Thermal Performance Curve fits for all species: Development Rate

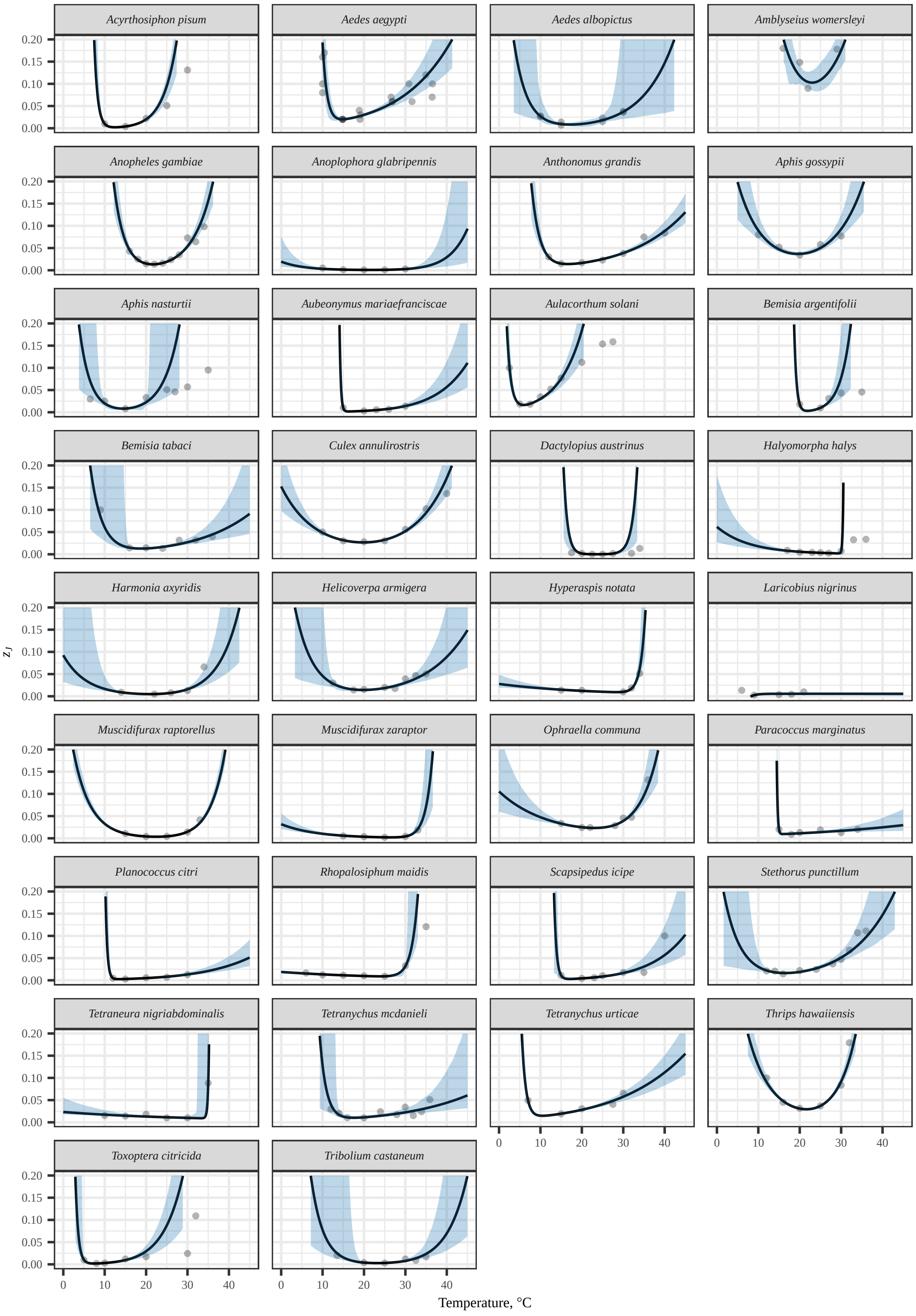

**Figure 8:** Thermal Performance Curve fits for all species: Juvenile Mortality

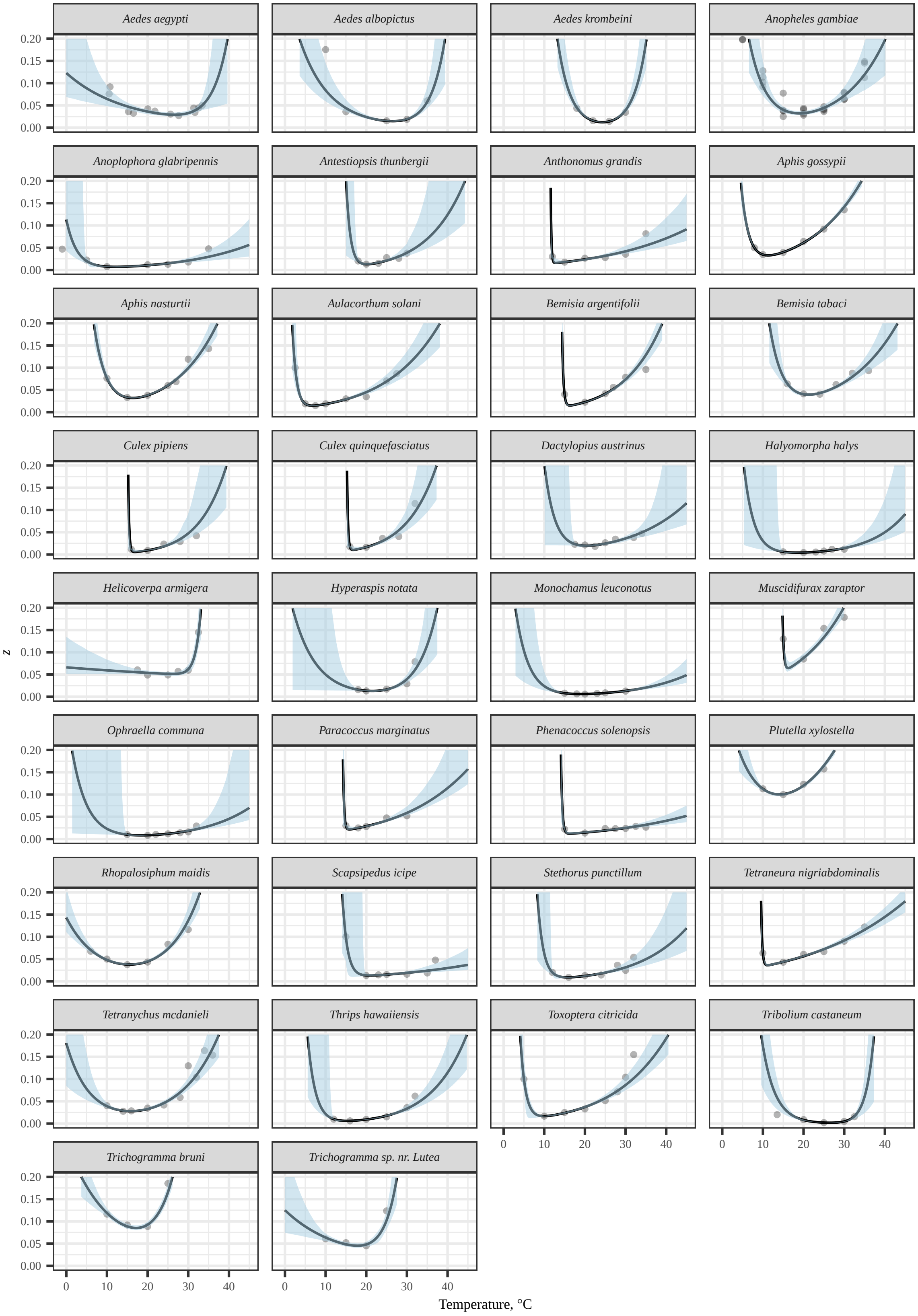

**Figure 9:** Thermal Performance Curve fits for all species: Adult Mortality

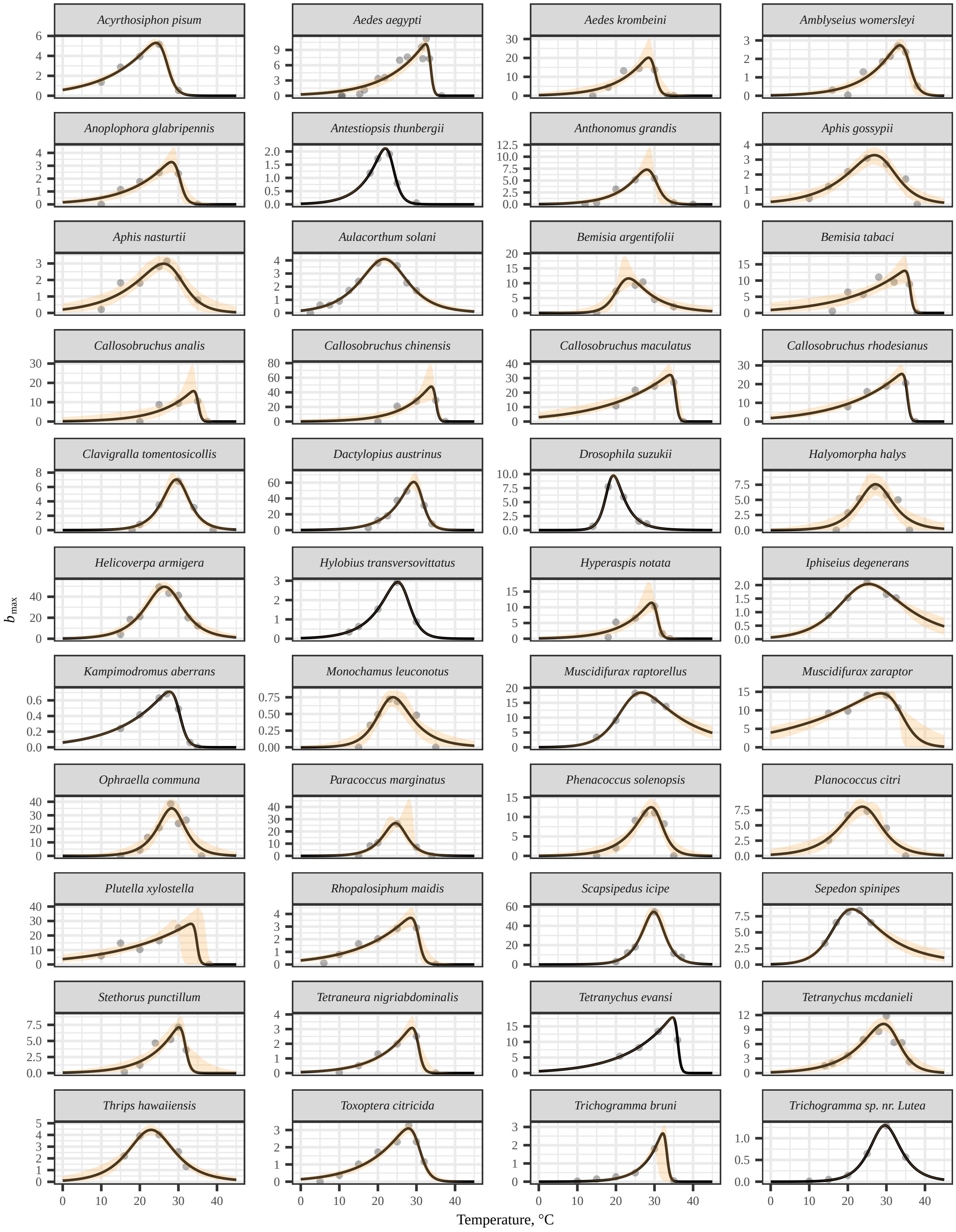

**Figure 10:** Thermal Performance Curve fits for all species: Fecundity

#### 1.6 Species-level temperature dependencies of $r_m$

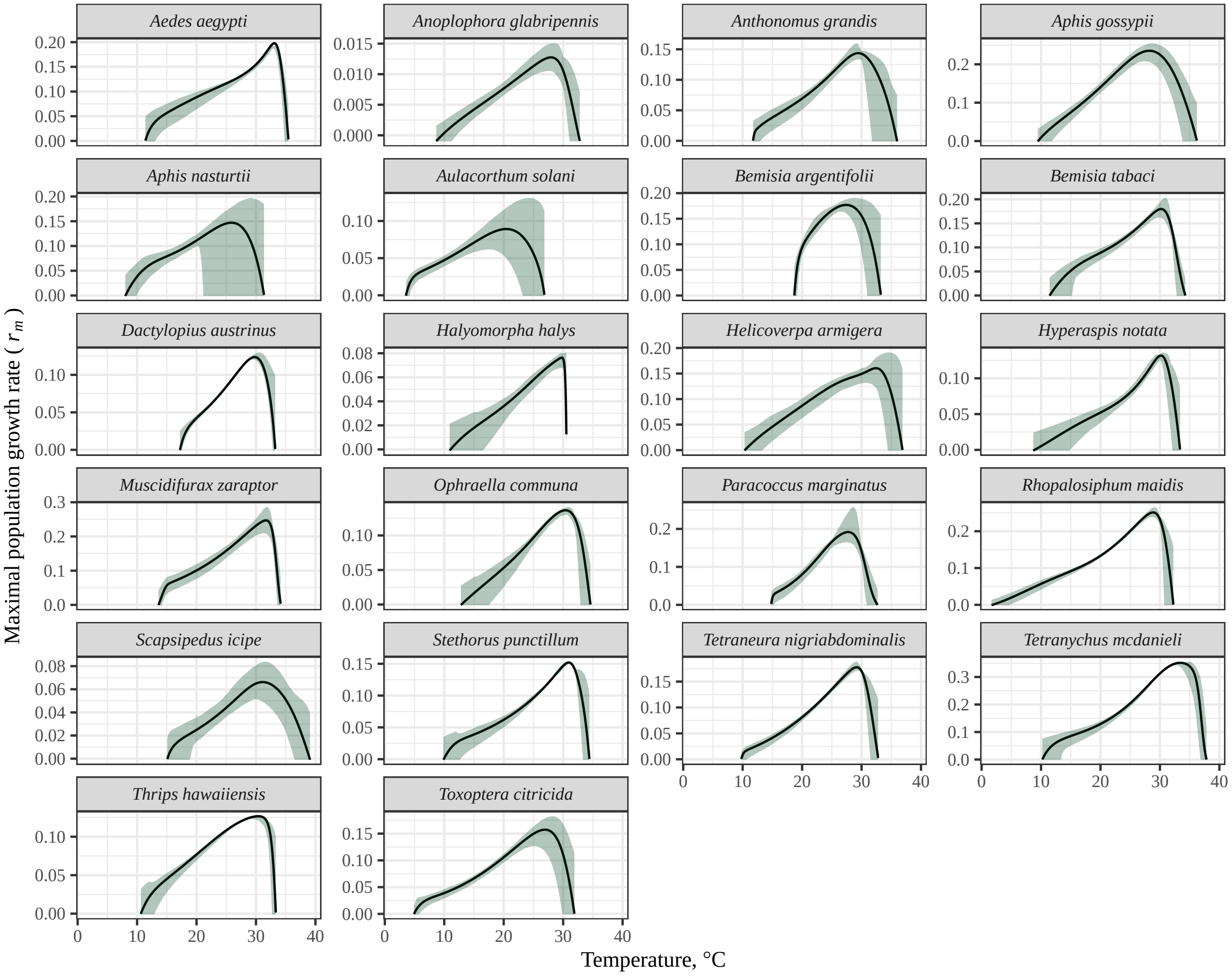

**Figure 11:** Thermal Performance Curve fits for all species: $r_m$

#### 1.7 Sensitivity of the results to the parameterisation of fecundity loss rate ($\kappa$)

Fecundity typically declines over time, which can have significant impacts on the lifetime reproduction of individuals and therefore fitness. The rate at which fecundity declines with age ($\kappa$) may be temperature-dependent, but there appears to be practically no existing data on this for arthropods. Therefore, here we quantify the sensitivity of our theoretical predictions to changes in parameterisation of baseline fecundity loss rate (the normalisation constant, $\kappa_0$). Specifically, we re-evaluate our trait sensitivity analyses, as well as our calculation of selection gradient by varying $\kappa_0$ across two extreme values, around the value we have used to generate the main results (0.1). Supplementary Figure 12 shows how changing $\kappa_0$ affects the shape of the fecundity curve at any given temperature.

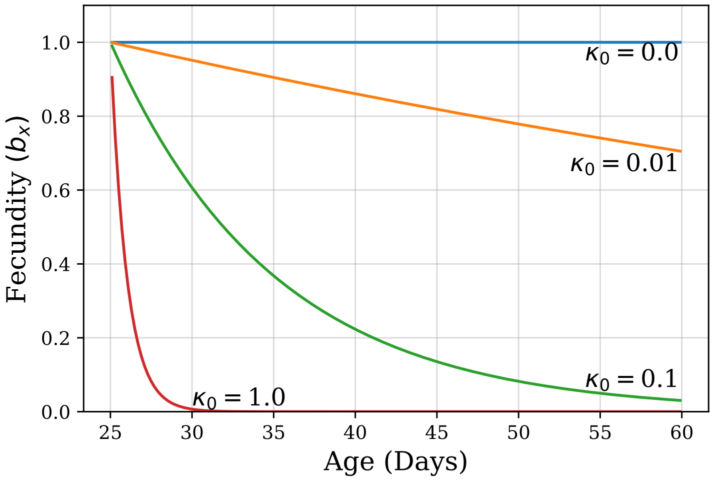

**Figure 12:** Sensitivity of the fecundity TPC to changes in $\kappa_0$.

Supplementary Figure 13 shows that the TPC shape for $\kappa$ remains qualitatively the same (for the two meaningful extreme values of $\kappa_0$):

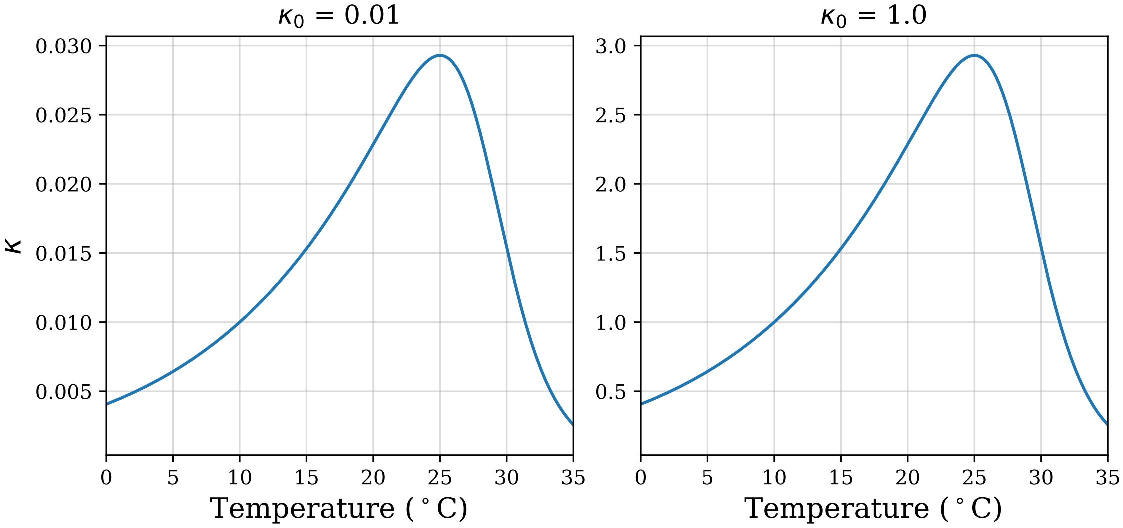

**Figure 13:** Insensitivity of the $\kappa$ curve to meaningfully extreme $\kappa_0$ values.

##### 1.7.1 Effect on the trait sensitivity results

First, we re-evaluate the trait sensitivity analysis results (SM Fig. 14). As expected, in the case where baseline kappa ($\kappa_0$) is lower, maximum fecundity ($b_{max}$) becomes more important relative to $\kappa$, leaving the order of importance of the 5 traits the same as for the intermediate case ($\kappa_0$=0.1) upon which our main results are based.

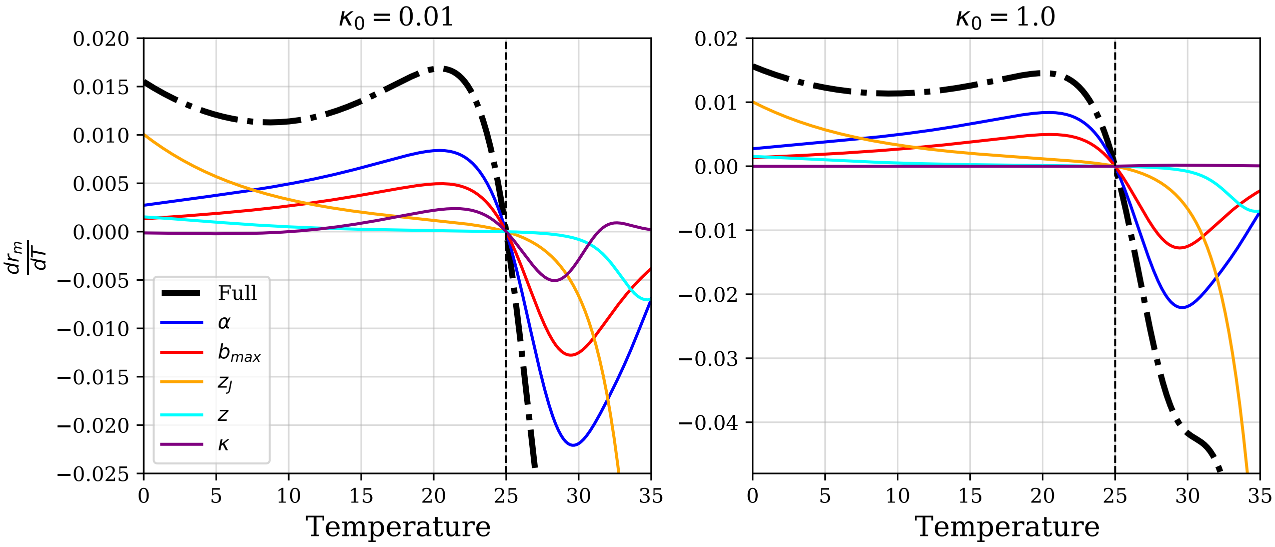

**Figure 14:** Re-evaluation of the results of the trait sensitivity analysis

##### 1.7.2 The selection gradients revisited

Next we re-evaluate the $r_m$ TPC and selection gradients as above. We focus only on the dominant trait $\alpha$ because the order of the strengths of selection gradients is bound to remain unchanged due to the unchanged order of trait sensitivity irrespective of the $\kappa_0$ value (previous section). Supplementary Figure 15 shows that the selection gradient remains qualitatively unchanged, with overall $r_m$ lower when $\kappa_0$ is high, as expected.

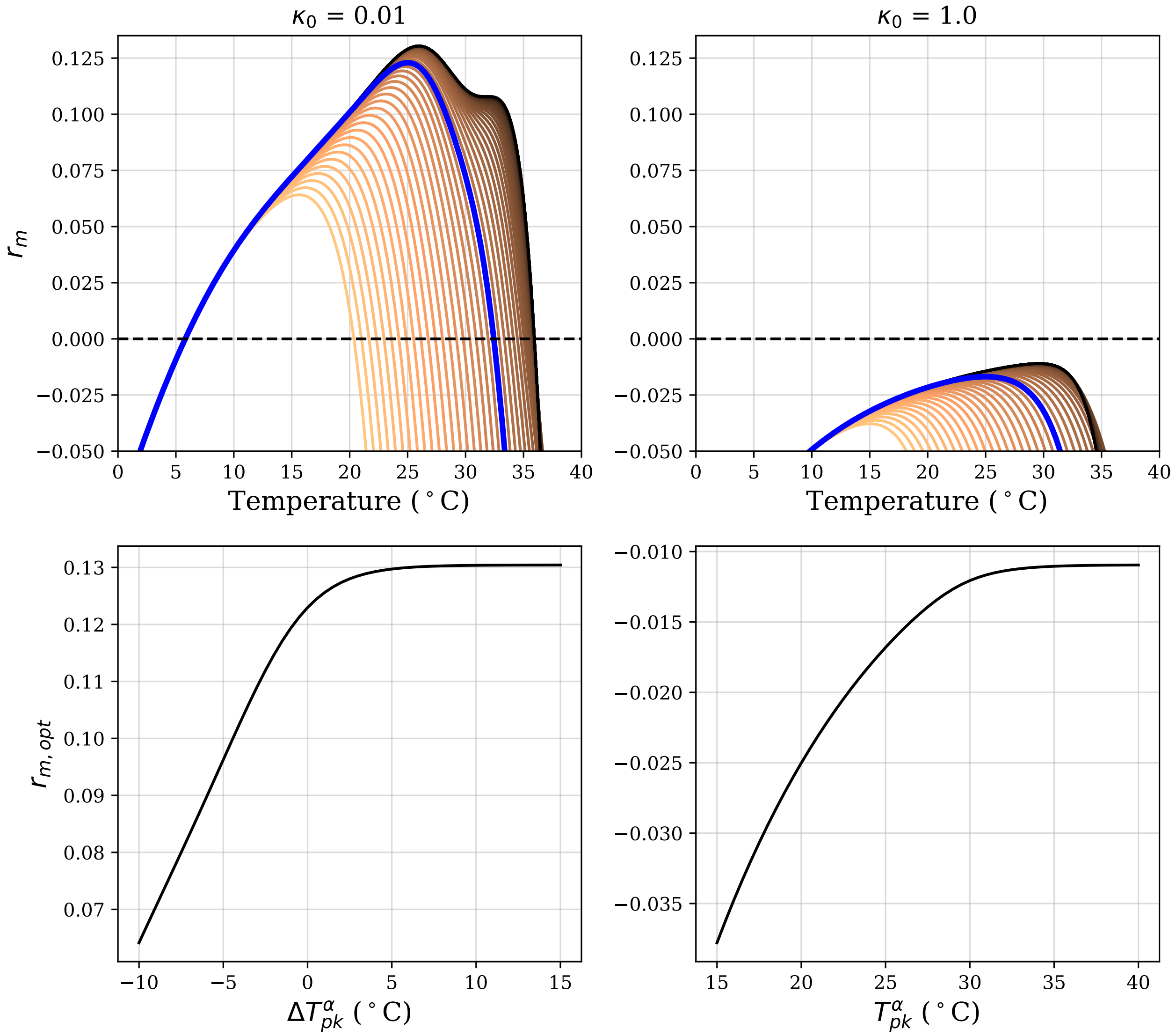

**Figure 15:** Insensitivity of $r_m$ selection gradient to changes in $\kappa_0$

#### 1.8 Macroevolutionary patterns and phylogenetic constraints

**Table 1:** Nucleotide sequences collected for each species from the SILVA (SSU and LSU) and Barcode of Life Data System (COI-5P) databases.

| **Species** | **SSU ID** | **LSU ID** | **COI-5P ID** |
|---|---|---|---|
| *Acyrthosiphon pisum* | ABLF02002530.1.2075 | F02004049.1.2297 | - |
| *Aedes aegypti* | AAGE02033765.6222.8206 | AAGE02025420.70.4046 | CULSA016-19 |
| *Aedes albopictus* | GCLM01041991.604.2545 | MNAF02000533.14119.17736 | ACMIP154-07 |
| *Aedes krombeini* | - | - | - |
| *Amblyseius swirskii* | - | - | GBMNC68842-20 |
| *Amblyseius womersleyi* | - | - | GBCH5643-13 |
| *Anopheles gambiae* | AM157179.1.2015 | LCWJ01002898.1.3298 | CULSA066-19 |
| *Anoplophora glabripennis* | - | - | GBMNE15612-21 |
| *Antestiopsis thunbergii* | - | - | - |
| *Anthonomus grandis* | EU215423.9073.11005 | EU215423.12455.16248 | GBMIN12012-13 |
| *Aphis gossypii* | - | - | ACEA143-14 |
| *Aphis nasturtii* | - | - | ACEA530-14 |
| *Aubeonymus mariaefranciscae* | - | - | - |
| *Aulacorthum solani* | AF487713.1.966 | - | ACEA131-14 |
| *Bemisia argentifolii* | - | - | - |
| *Bemisia tabaci* | GCZW01018184.195.2694 | - | BTB002-12 |
| *Callosobruchus analis* | - | - | GBCCH430-13 |
| *Callosobruchus chinensis* | - | - | CSP030-09 |
| *Callosobruchus maculatus* | GEUE01061439.889.2802 | GEUD01155100.99.2572 | GBCCH431-13 |
| *Callosobruchus rhodesianus* | - | - | GBCL2570-06 |
| *Clavigralla tomentosicollis* | GAJX01000019.410.2325 | - | GBMNA17782-19 |
| *Culex annulirostris* | - | - | GBMNC791-20 |
| *Culex pipiens* | AY988445.1.1858 | - | CULSA020-19 |
| *Culex quinquefasciatus* | AAWU01003351.41983.43837 | AAWU01047416.6583.8759 | GBDP12712-12 |
| *Dactylopius austrinus* | AY795538.1.608 | - | - |
| *Drosophila suzukii* | AWUT01011932.32096.34063 | AWUT01017126.2.3394 | GBDPD245-14 |
| *Gastrolina depressa* | - | - | GBMNA17939-19 |
| *Halyomorpha halys* | GEDY01000115.149.2061 | GBHT01004590.7.2044 | AGIRI147-17 |
| *Harmonia axyridis* | KP419116.1.1832 | - | GBCL17655-14 |
| *Helicoverpa armigera* | KT343378.1.1903 | - | GBGL29849-19 |
| *Hylobius transversovittatus* | - | - | GBCL3480-08 |
| *Hyperaspis notata* | - | - | HEAUG012-12 |
| *Iphiseius degenerans* | - | - | TZBCA378-07 |
| *Kampimodromus aberrans* | - | - | - |
| *Laricobius nigrinus* | KP419143.1.1857 | - | GBCL9856-12 |
| *Leptinotarsa decemlineata* | GEEF01054810.5881.7790 | - | FBCOP312-13 |
| *Macrocentrus iridescens* | - | - | BBHEC285-09 |
| *Macrosiphum euphorbiae* | - | - | ACEA250-14 |
| *Monochamus leuconotus* | - | - | - |
| *Muscidifurax raptorellus* | - | - | - |
| *Muscidifurax zaraptor* | - | - | - |
| *Myzus persicae* | LXJY01000320.20922.22509 | - | GBMNE22299-21 |
| *Ophraella communa* | - | - | GBCCH11138-19 |
| *Otiorhynchus sulcatus* | AF250084.1.1795 | - | COLFD838-12 |
| *Paracoccus marginatus* | FIZT01020513.6507.8206 | - | GBMIN46559-16 |
| *Phenacoccus solenopsis* | - | - | GBMHH29894-19 |
| *Planococcus citri* | GAXF02029206.892.3340 | - | GBMIN46549-16 |
| *Plutella xylostella* | AHIO01004014.23148.25041 | - | AACTA1852-20 |
| *Rhopalosiphum maidis* | - | - | ACEA794-14 |
| *Scapsipedus icipe* | - | - | GBMOR7430-19 |
| *Sepedon spinipes* | - | - | FIDIP848-12 |
| *Stethorus punctillum* | EF512328.1.1788 | - | ASCMT030-11 |
| *Tetraneura nigriabdominalis* | - | - | ASHMT220-11 |
| *Tetranychus evansi* | AB926295.1.1858 | - | GACAC126-12 |
| *Tetranychus mcdanieli* | - | - | - |
| *Tetranychus urticae* | CAEY01001788.15245.16831 | AY750693.1.2826 | GACAC6465-19 |
| *Thrips hawaiiensis* | - | - | AGIMP052-16 |
| *Toxoptera citricida* | AY216697.1.2480 | - | RDBA411-06 |
| *Tribolium castaneum* | HM156711.1.1831 | - | BIPR011-13 |
| *Trichogramma bruni* | - | - | - |
| *Trichogramma* sp. nr. *Lutea* | - | - | KMPUH785-19 |

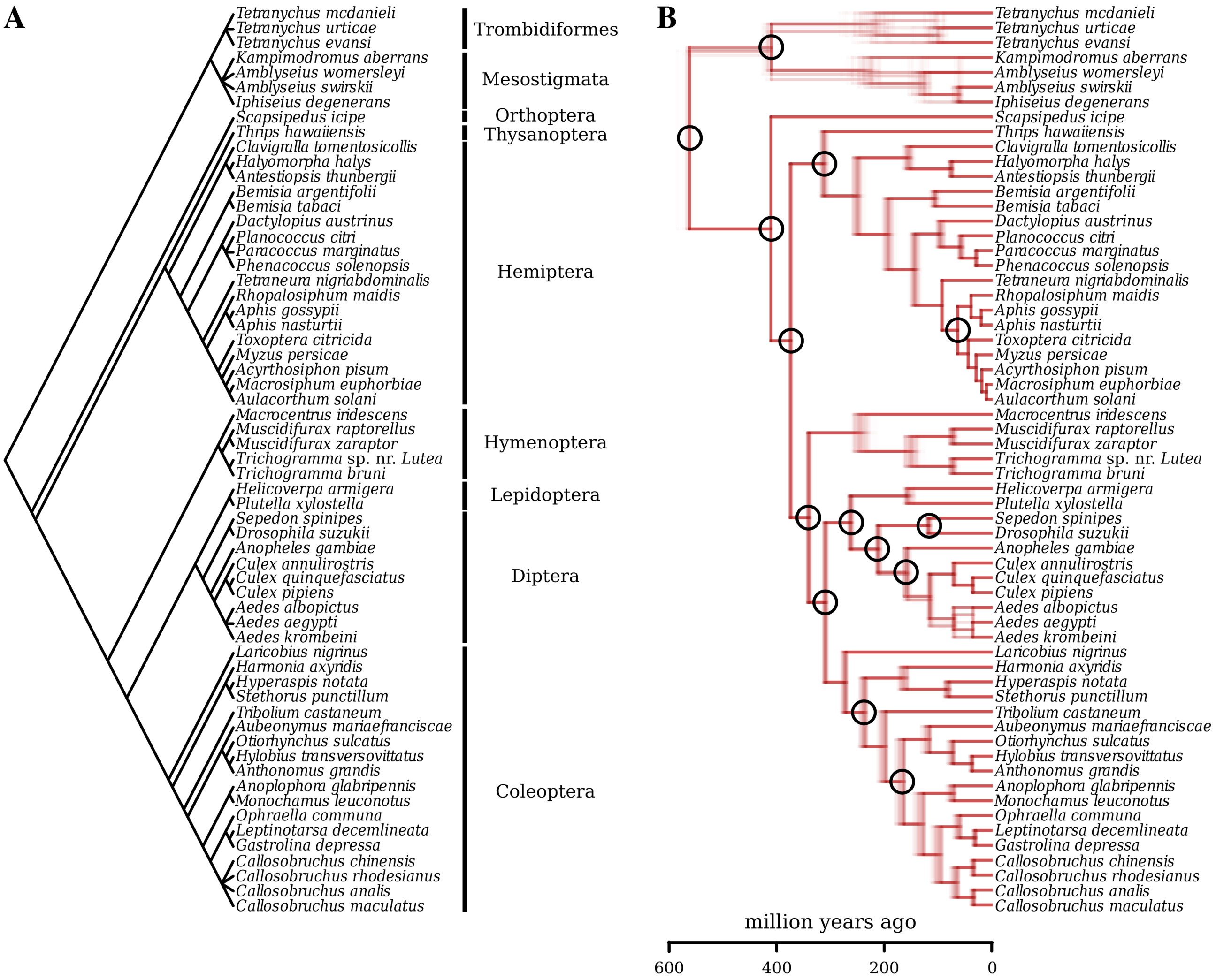

**Figure 16:** A: The Open Tree of Life topology for the species in the study, with orders explicitly shown. Note that this topology includes some polytomies (e.g., see the *Callosobruchus* and *Tetranychus* clades). B: The final set of 100 time-calibrated trees overlaid on top of each other. Differences in both topology and branch lengths can be observed. Some polytomies from panel A were objectively resolved (e.g., the *Callosobruchus* clade) based on the concatenated sequence alignment. In contrast, where sequence data were completely missing for at least one species (e.g., the *Tetranychus* clade), polytomies were randomly resolved. Nodes whose age was obtained from the TimeTree database are marked with a circle. The two panels were plotted with the ape [5] (v.5.6-2) and phytools R packages [6] (v.1.2-0), respectively.

### References

[1] Savage, V. M., Deeds, E. J. & Fontana, W. Sizing up allometric scaling theory. *PLoS computational biology* **4**, e1000171 (2008).

[2] Kontopoulos, D.-G. *et al.* Phytoplankton thermal responses adapt in the absence of hard thermodynamic constraints. *Evolution* **74**, 775–790 (2020).

[3] Frazier, M., Huey, R. B. & Berrigan, D. Thermodynamics constrains the evolution of insect population growth rates:"warmer is better". *Am. Nat.* **168**, 512–520 (2006).

[4] Pawar, S., Dell, A. I., Savage, V. M. & Knies, J. L. Real versus Artificial Variation in the Thermal Sensitivity of Biological Traits. *Am. Nat.* **187**, E41–E52 (2016).

[5] Paradis, E. & Schliep, K. ape 5.0: an environment for modern phylogenetics and evolutionary analyses in R. *Bioinformatics* **35**, 526–528 (2019).

[6] Revell, L. J. phytools: an R package for phylogenetic comparative biology (and other things). *Methods in Ecology and Evolution* **3**, 217–223 (2012).
